# Convergent evolution of plant prickles is driven by repeated gene co-option over deep time

**DOI:** 10.1101/2024.02.21.581474

**Authors:** James W. Satterlee, David Alonso, Pietro Gramazio, Katharine M. Jenike, Jia He, Andrea Arrones, Gloria Villanueva, Mariola Plazas, Srividya Ramakrishnan, Matthias Benoit, Iacopo Gentile, Anat Hendelman, Hagai Shohat, Blaine Fitzgerald, Gina M. Robitaille, Yumi Green, Kerry Swartwood, Michael J. Passalacqua, Edeline Gagnon, Rebecca Hilgenhof, Trevis D. Huggins, Georgia C. Eizenga, Amit Gur, Twan Rutten, Nils Stein, Shengrui Yao, Clement Bellot, Mohammed Bendahmane, Amy Frary, Sandra Knapp, Tiina Särkinen, Jesse Gillis, Joyce Van Eck, Michael C. Schatz, Yuval Eshed, Jaime Prohens, Santiago Vilanova, Zachary B. Lippman

## Abstract

An enduring question in evolutionary biology concerns the degree to which episodes of convergent trait evolution depend on the same genetic programs, particularly over long timescales. Here we genetically dissected repeated origins and losses of prickles, sharp epidermal projections, that convergently evolved in numerous plant lineages. Mutations in a cytokinin hormone biosynthetic gene caused at least 16 independent losses of prickles in eggplants and wild relatives in the genus *Solanum*. Strikingly, homologs promote prickle formation across angiosperms that collectively diverged over 150 million years ago. By developing new *Solanum* genetic systems, we leveraged this discovery to eliminate prickles in a wild species and an indigenously foraged berry. Our findings implicate a shared hormone-activation genetic program underlying evolutionarily widespread and recurrent instances of plant morphological innovation.

## Introduction

Trait convergence, defined as the emergence of analogous traits in distantly related organisms, was a key observation made by Darwin in support of his theory of evolution. He recognized that similar selective pressures could lead to similar yet independently derived adaptations across species. However, the extent to which phenotypic convergence is driven by corresponding convergence in underlying genetic programs is poorly understood. Within a species, adaptive traits may arise from selection acting on standing genetic variation within and among populations, making phenotype-genotype convergence more likely (*1, 2*). At higher taxonomic levels and with increasing evolutionary divergence, phenotype-genotype convergence is posited to decline due to variation in allelic diversity, genomic background, and developmental mechanisms (*3, 4*). However, opportunities to dissect convergence at these timescales are scarce; finding convergent traits across wide evolutionary spans that are genetically tractable and well-supported by genomic data has remained a significant challenge.

In plants, sharp epidermal projections known as prickles convergently evolved at least 28 times over more than 400 million years of evolution (**Fig. 1A and Table S1**). Prickles serve adaptive functions in herbivore deterrence, climbing growth, plant competition, and water retention (*5–8*). Rose (*Rosa* spp.) is a widely recognized taxon bearing prickles, though these prickles are vernacularly called thorns. True thorns, which are found on the trees of citrus (*Citrus* spp.) and honey locusts (*Gleditsia* spp.), for example, develop from axillary branches, whereas prickles originate from the epidermis or cortex, typically in association with hair-like structures known as trichomes (*5*). Despite their diverse adaptive roles and the broad phylogenetic diversity of their origins, prickles exhibit remarkable morphological similarity (**Fig. S1A-C**). Moreover, prickles have been lost or suppressed in numerous lineages. Therefore, prickle formation is an attractive system to determine whether episodes of repeated trait evolution rely on the same genetic programs over both short and long evolutionary timescales.

**Fig. 1.**
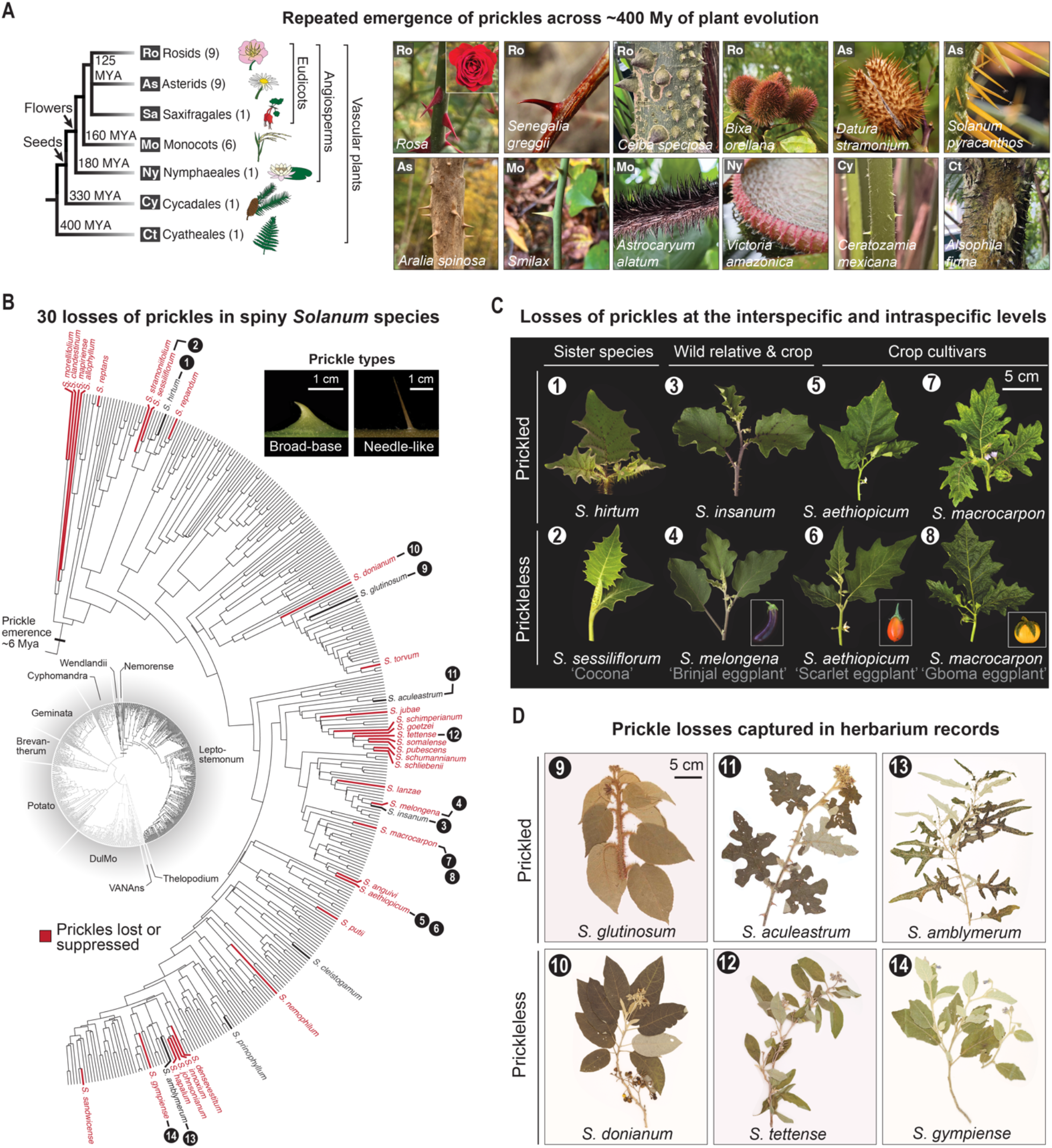
Prickles evolved convergently across vascular plants and were lost repeatedly in the spiny *Solanum* lineage. (**A**) Phylogeny (reproduced from https://lifemap-ncbi.univ-lyon1.fr/) and corresponding images of representative vascular plants that independently evolved prickles. Number in parentheses indicates number of identified independent evolutionary origins of prickles (**B**) Phylogenetic tree [adapted from (*9, 10*)] of the spiny *Solanum* (subclades Wendlandii, Nemorense, and Leptostemonum) with species having lost prickles highlighted in red. Representative images of narrow and broad-based prickle morphologies are shown. (**C** and **D**) Images of *Solanum* taxa that have lost prickles captured from living (C) and herbarium (D) collections. Numbers correspond to species shown in (B).

In the genus *Solanum*, which includes the major crops eggplant, potato, and tomato, prickles emerged in the common ancestor of the so-called “spiny *Solanums*” around 6 million years ago (Mya) (*9, 10*). This lineage includes the large Leptostemonum clade, which comprises hundreds of globally distributed species, including all cultivated eggplants and their wild progenitors (**Fig. 1B**). Prickle morphologies across the clade range from broad at the base (broad-based), or narrow-based and needle-like. Prickles occur on stems, along the vasculature of leaves, and on calyces, the outer whorl of floral organs. Several spiny *Solanum* species underwent human-driven selection for losses or suppression of prickles, facilitating comparisons of prickled and prickleless sister species, crop species, and wild relatives (**Fig. 1C and Table S2**). An agriculturally significant instance of prickle loss occurred during the domestication of the widely cultivated Brinjal eggplant (*S. melongena*); however, prickle losses have also been observed in wild *Solanum* species without history of domestication (**Fig. 1D**). The specific genes controlling prickle development are unknown.

## Results

### Repeated losses of prickles in cultivated eggplants are caused by *LOG* gene mutations

Previous mapping studies in Brinjal eggplant showed that the loss of prickles is inherited as a single Mendelian locus designated *prickleless* (*pl*) and localized to a genomic interval on chromosome 6 (*11*). Using a recurrent backcross-derived mapping population between Brinjal eggplant and its prickled wild progenitor *S. insanum*, we confirmed this result and further fine-mapped *pl* to a ∼100 kb interval containing 10 annotated genes (**Fig 2A**). Just outside this interval is the previously proposed *pl* candidate gene *SmelARF18*, a putative auxin hormone response transcription factor (*12*). However, we did not find conspicuous coding region loss-of-function mutations in this gene or in any other gene in the interval. Instead, we identified a probable splice-site mutation in a gene encoding a LONELY GUY (LOG)-family cytokinin biosynthetic enzyme. LOG family members catalyze the final step in the biosynthesis of bioactive cytokinin, a hormone with roles in plant cell proliferation and differentiation (*13*). In a collection of 23 re-sequenced eggplant accessions, we found that this splice-site mutation was consistently associated with the prickleless phenotype, except in one accession, which harbored a 474 bp deletion in exon 6 of the *LOG* gene (**Fig. S2A**).

**Fig. 2.**
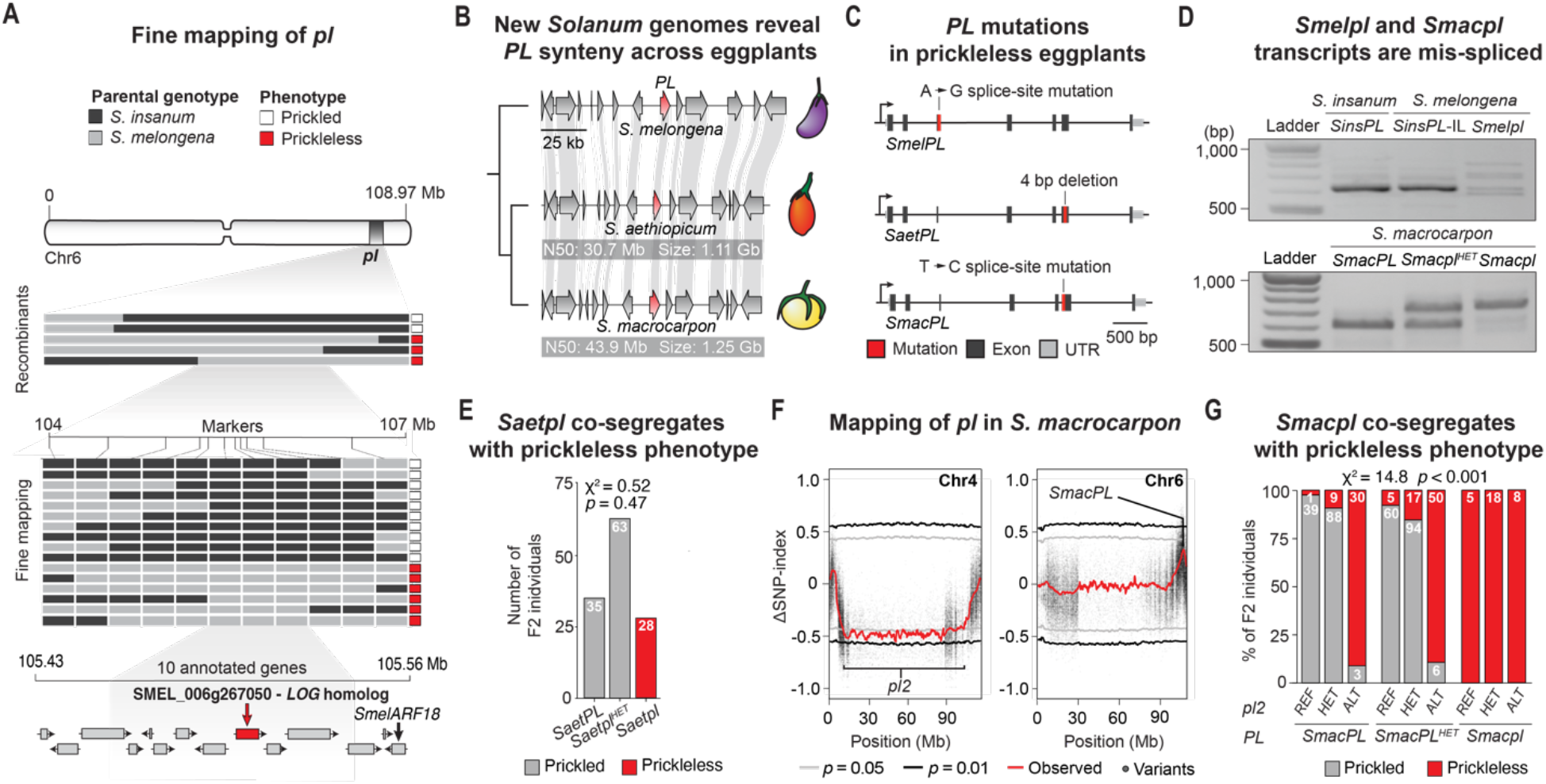
Losses of prickles in three domesticated *Solanum* species are caused by independent mutations in a *LOG* cytokinin biosynthetic gene. (**A**) Fine-mapping of *pl* in a Brinjal eggplant (*S. melongena*) x wild progenitor species (*S. insanum*) mapping population. (**B**) Genome sequencing and chromosome-scale assemblies of two African eggplants, the Scarlet eggplant (*S. aethiopicum*) and the Gboma eggplant (*S. macrocarpon*) reveals synteny of the *pl* locus. Genome summary statistics are indicated. (**C**) Independent mutations in a *LOG* gene in the *pl* interval in all three prickleless crop species. (**D**) Mis-splicing of *PL* transcripts caused by the *pl* mutations in Bringal eggplant *pl (Smelpl*) and Gboma eggplant *pl (Smacpl*) confirmed by RT-PCR. *SinsPL*-IL denotes an introgression of *S. insanum PL* into the Brinjal eggplant genomic background. (**E**) *Saetpl* co-segregates with the prickleless phenotype in a Scarlet eggplant F2 mapping population. (**F**) QTL-Seq identifies two loci that independently cause the prickleless phenotype in Gboma eggplant. (**G**) F2 population confirms co-segregation of *Smacpl* and *pl2* with the prickleless phenotype. Numbers within bars (E and G) indicate count totals of individuals within each genotype class.

The discovery of two independent mutations in the *LOG* candidate gene suggested that the loss of prickles occurred at least twice during Brinjal eggplant’s domestication history. It also raised the possibility that mutations in orthologous genes may have caused parallel prickle losses in two other independently domesticated African eggplant species, the Scarlet eggplant (*S. aethiopicum*) and the Gboma eggplant (*S. macrocarpon*). Genomic resources for these indigenous crop species are limited. We therefore sequenced and assembled high-quality (QV ≥ 51, completeness > 99) chromosome-scale genomes and generated gene annotations for both species (**Fig. 1C, Tables S3 and S4**). Using these resources, we found that synteny within the *pl* locus was retained across all three cultivated eggplant species (**Fig. 2B**), and that prickleless Scarlet eggplant and Gboma eggplant each harbored different loss-of-function mutations in their respective *LOG* orthologs (**Fig. 2C**). Scarlet eggplant carries an indel mutation leading to a frameshift in the coding sequence and a prematurely terminated protein product, while Gboma eggplant carries a splice-site mutation. Reverse-transcriptase polymerase chain reaction (RT-PCR) on cDNA revealed lower expression and multiple mis-spliced transcripts in Brinjal eggplant and a mis-spliced isoform with a retained intron in Gboma eggplant (**Fig. 2D**). PCR sequencing revealed these transcripts were non-functional (**Fig. S2B-D**).

To further validate that these independent mutations explain the prickleless phenotypes we performed co-segregation analysis in F2 populations derived from intraspecific crosses between prickled and prickleless parents (**Fig. 1C and 2E**). In Scarlet eggplant, homozygosity of the *LOG* mutant allele co-segregated with the prickleless phenotype in a Mendelian recessive fashion in all examined individuals (χ^2^ = 0.52, df = 1, *p* = 0.47). Interestingly, in Gboma eggplant we observed segregation patterns that indicated the presence of another unlinked recessive variant independently contributing to prickle loss (χ^2^ = 14.8, df = 1, *p* < 0.001). Leveraging our newly developed genomic resources, we used a mapping-by-sequencing approach to identify a second large interval associated with the loss of prickles on chromosome 4, which we designated *pl2* (**Fig. 2F**). Importantly, all segregating homozygous mutant individuals at *pl* on chromosome 6 carried the *LOG* gene splice-site mutation, although this genotype class was represented at lower-than-expected frequency, likely owing to segregation distortion (**Fig. 2G**). Taken together, these results indicate that *PL* is the *LOG* candidate gene, and that at least four independent mutations in this gene enabled repeated selection for losses of prickles in cultivated eggplant species.

### Mutations in *PL* are found in prickleless wild and cultivated species across the *Solanum*

The clade encompassing all three of the cultivated eggplants diverged ∼2 Mya, but prickles in *Solanum* are more ancient, having emerged over ∼6 Mya, and 30 independent losses of prickles have been documented, including in additional domesticated and wild species (*10*). We tested whether mutations in *PL* underlie these repeated instances of prickle loss across this broader evolutionary timescale by sampling DNA from additional prickleless species and their prickled close relatives. Because many wild *Solanum* species are too rare or geographically inaccessible for live-tissue sampling, we used a combination of PCR-amplified exon sequencing from herbarium tissue samples and whole-gene sequencing from available live tissue samples to detect *PL* mutations (**Table S5**).

Along with the four *PL* mutations identified in our analysis of prickleless eggplants, we identified an additional 12 allelic mutations predicted to deleteriously affect *PL* function at the pan-genus level across the spiny *Solanum* (**Fig. 3**). These mutations explained 14 out of the 30 recorded losses of prickles across the genus, spanning species within and across subclades (**Fig. 1B and Table S6**). In some cases, we detected the same, although not necessarily ancestrally derived, alleles in separate species. For example, the same splice-site mutation found in prickleless Gboma eggplant, native to and cultivated almost exclusively in Africa, was also identified in the wild species *S. donianum*, whose native range is in Central America and the Caribbean. Likewise, an identical splice-site mutation was found in both the wild species *S. lanzae*, from western Africa, and the foraged and sometimes cultivated species *S. stramoniifolium*, native to northern South America. Such genetic convergence at the allelic level may reflect the high penetrance of *PL* splicing defects, which can be conferred by mutationally accessible single nucleotide variants (*14*). Together, our results suggest that *PL* had an important and repeated genetic role in the convergent losses of prickles across *Solanum* in the wild and in cultivation.

**Fig. 3.**
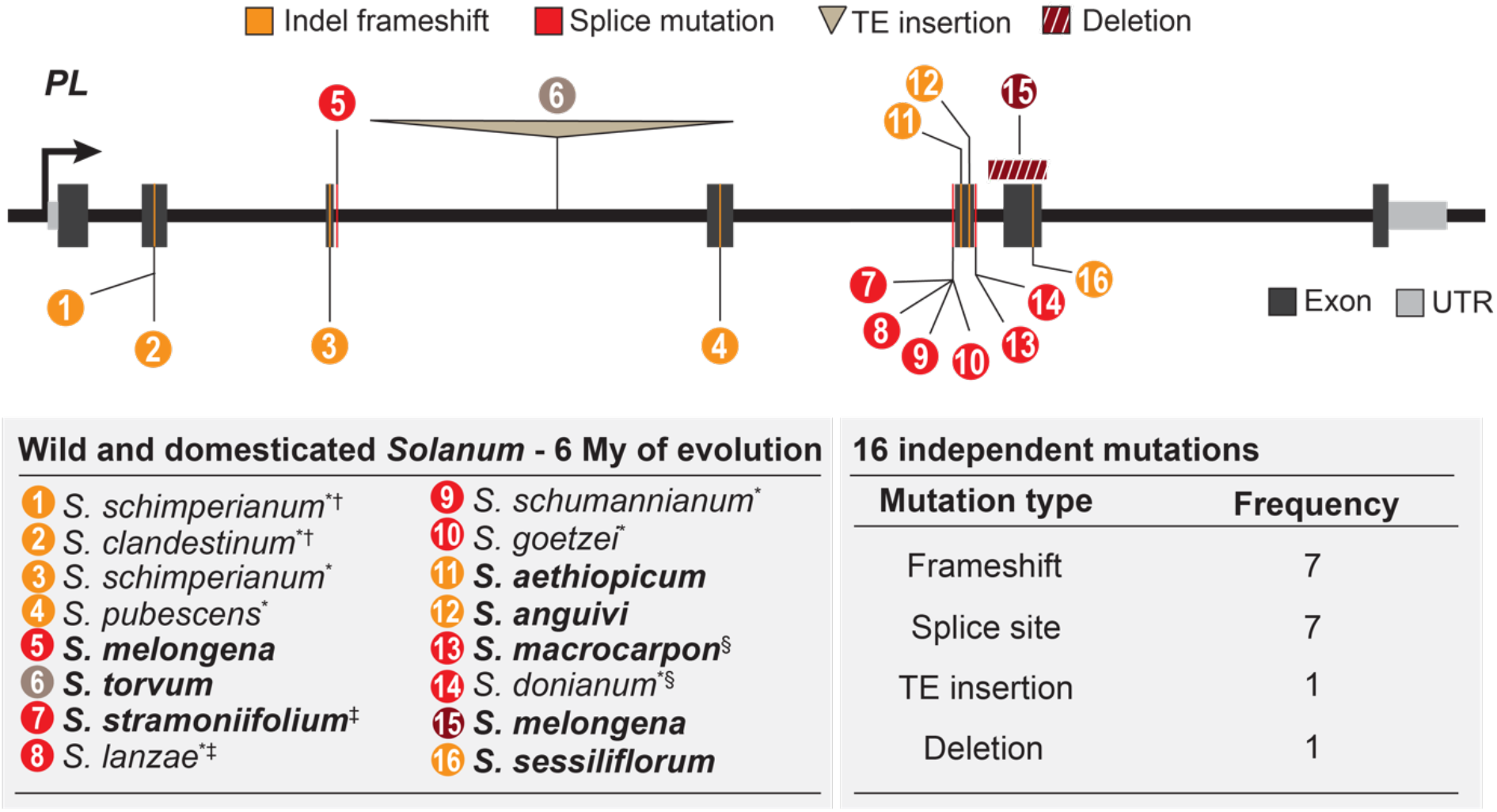
Mutations in *PL* are associated with prickle suppression across the spiny *Solanum*. *PL* variants with strong probable deleterious effects on gene function identified in prickle-suppressed taxa but not in closely-related prickled sister taxa. Mutations are numbered and shown along with their corresponding species name and sample source in the table below. In the tables, bold text indicates cultivated species, (*) indicates that genotyping was performed on archival herbarium samples, (†,‡,§) indicate species pairs that share identical but not necessarily ancestrally-derived mutations.

### Repeated co-option of *LOG* homologs underlies prickle convergent evolution

The finding of recurrent mutations in *PL* orthologs across the spiny *Solanums* suggested that co-option of cytokinin biosynthetic gene function was critical to prickle evolution. This spurred us to ask whether genetic convergence through *LOG* gene co-option extends to other prickled species across flowering plants. We searched the literature for studies associating instances of loss or suppression of sharp outgrowths with specific genomic loci or genes. Strikingly, we found that in the grass family (Poaceae) independent mutant alleles in a *LOG* homolog from rice (*Oryza sativa*) and barley (*Hordeum vulgare*) conferred near complete suppression of epidermally-derived sharp projections commonly called “barbs” but botanically classified as prickles (*15, 16*). In contrast to the conspicuous, multi-cellular, and lignified prickles found in the *Solanum* (**Fig. S1**), grass prickles are homologous structures made of silicified single-cells that develop on awns, an outer-whorl structure of the grass flower involved in seed dispersal (**Fig 4A**).

**Fig. 4.**
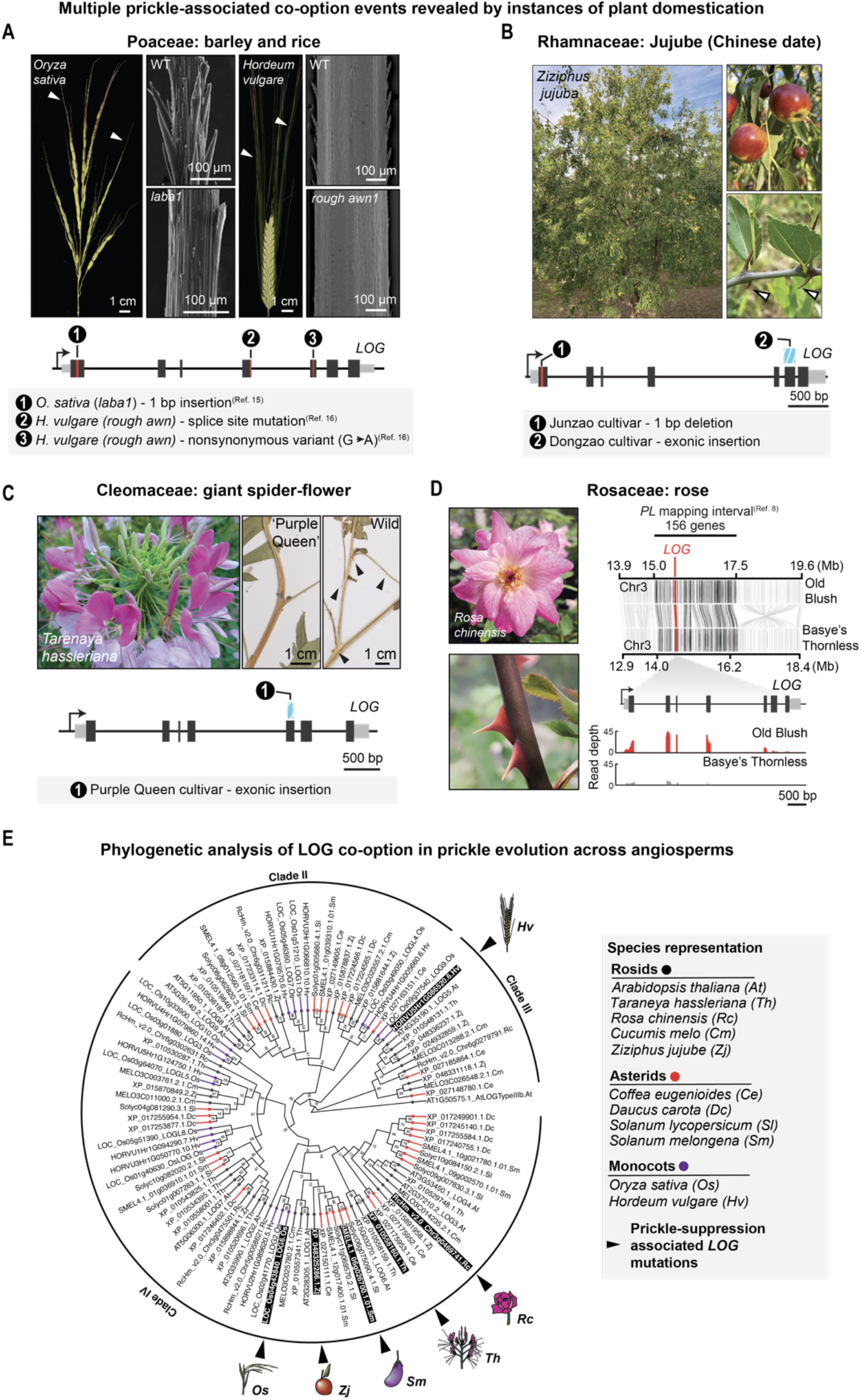
Losses of convergently evolved prickles across angiosperms are associated with *LOG* mutations. (**A** to **D**) Instances of prickle suppression in angiosperms associated with *LOG* mutations depicted in corresponding *LOG* gene diagrams. (A) Images of rice and barley WT inflorescences. Arrowheads indicate awns, which are shown for WT and mutant genotypes (rice, *laba1*; barley, *rough awn1*) by SEM. (B) Images of jujube trees, fruits, and stipular spines (arrowheads). Two less spiny cultivated varieties harbor two independent *LOG* mutations. (C) The ornamental giant spider flower (pictured) carries a mutated *LOG* gene in the sequenced ‘Purple Queen’ cultivar. Cultivated varieties bear fewer smaller prickles (arrowheads) than wild varieties, as reflected in herbarium samples (D). Loss of prickles in rose maps to a ∼2.5 Mb interval harboring a *LOG* gene with severely reduced expression in the prickleless cultivar relative to the prickled cultivar. Syntenic genes within the mapping interval of the prickled ‘Old Blush’ and prickleless ‘Basye’s Thornless’ parental lines are shown in black. Read pileups show average *LOG* expression in leaves of the parental genotypes (*n* = 3). (**E**) Protein-based phylogenetic tree of the Arabidopsis LOG1 orthogroup defined by Orthofinder, from the indicated asterid (red), rosid (black), and monocot (purple) species. LOGs encoded by genes with mutations in prickle-suppressed taxa are indicated by arrowheads.

Mining additional genomic data for *LOG* mutations co-occurring with losses of prickles in other eudicot lineages, we found that the fruit-bearing tree crop jujube, commonly known as Chinese date (*Ziziphus jujuba*), in the Rhamnaceae family, carrie two independent mutations (a 1 bp deletion and an exonic insertion) in a *LOG* homolog in two cultivars with suppressed prickles (also known as stipular spines) (**Fig. 4B**) (*17*–*19*). Importantly, neither mutation was found in Sour jujube (*Z. jujuba* var. *spinosa*), the prickled wild progenitor. We also detected an exonic insertion in a *LOG* homolog of the prickle suppressed ‘Purple Queen’ cultivar of giant spider flower (*Teranaya hassleriana*), a widely cultivated ornamental plant in the Cleomaceae, a small family within the Brassicales closely related to Arabidopsis (**Fig. 4C and Table S5**) (*20*). Finally, in rose, which is the most important cultivated cut flower, previous mapping for loci conferring “thornlessness” identified two major effect loci (*8*), as we found in *S. macrocarpon*. Notably, one of these was a ∼2.5 Mb interval containing 156 annotated genes (**Fig. 4D**), which we found includes a *LOG* homolog. Though there were no obvious coding or splicing mutations in this *LOG*, we found that its expression was substantially reduced in the leaves of the mapping parent cultivar ‘Bayse’s thornless’ (*Rosa wichuraiana*) compared to the prickled parent *Rosa chinensis* (**Fig. 4D**).

Taken together, these findings suggested that *LOG* gene reuse was critical in the independent acquisition of prickles in numerous plant lineages that last shared a common ancestor ∼150 Mya. Most sequenced seed plants (angiosperms and gymnosperms) retain multiple *LOG* gene copies within their genomes. In these taxa, the mean number of annotated *LOG* genes is 15, inflated by recent polyploid lineages, while the median and mode copy numbers are 12 and 10, respectively (*n* = 160). To understand the phylogenetic context of *LOG* co-option and to ask whether repeated co-option occurs in a specific clade of *LOG* gene family members, we conducted an analysis of LOG family proteins from prickled and prickleless species across the angiosperms (**Fig. 4E**). Most of the prickle co-option associated LOGs occurred within a specific subclade of the LOG family, suggesting that co-option was more favorable in certain LOG family subclades, particularly those with lineage-specific duplications. Intriguingly, however, the LOG homolog co-opted in barley is derived from an earlier diverging subclade (*16*), indicating that co-option of other LOG family members in different clades may also promote prickle evolution.

### *LOG* gene diversification preceded *PL* co-option in *Solanum*

Given the recurrent co-option of *LOG* genes against a backdrop of paralogous gene family members, we sought to better understand the phylogenetic and genomic context that facilitated *LOG* co-option in *Solanum*. We examined the conservation of the *PL* locus, comparing the region across *Solanum*, including Brinjal eggplant and two additional spiny *Solanum* species, with tomato (*S. lycopersicum*), an ancestrally prickleless species that diverged prior to the evolution of the spiny *Solanums*. We first constructed high-quality chromosome-scale genome assemblies for *S. prinophyllum* (Forest nightshade; QV = 51.6, completeness > 99) and *S. cleistogamum* [Desert raisin; QV = 49.8, completeness > 99 (**Tables S3 and S4**)]. Forest nightshade is endemic to southeastern Australia whereas Desert raisin is native to the arid center of Australia and has been foraged by First Nations people for thousands of years for their sweet, dried berries (*21*) (**Fig. 5A**). Notably, neither species has been domesticated and both are distinct lineages from the cultivated eggplants.

**Fig. 5.**
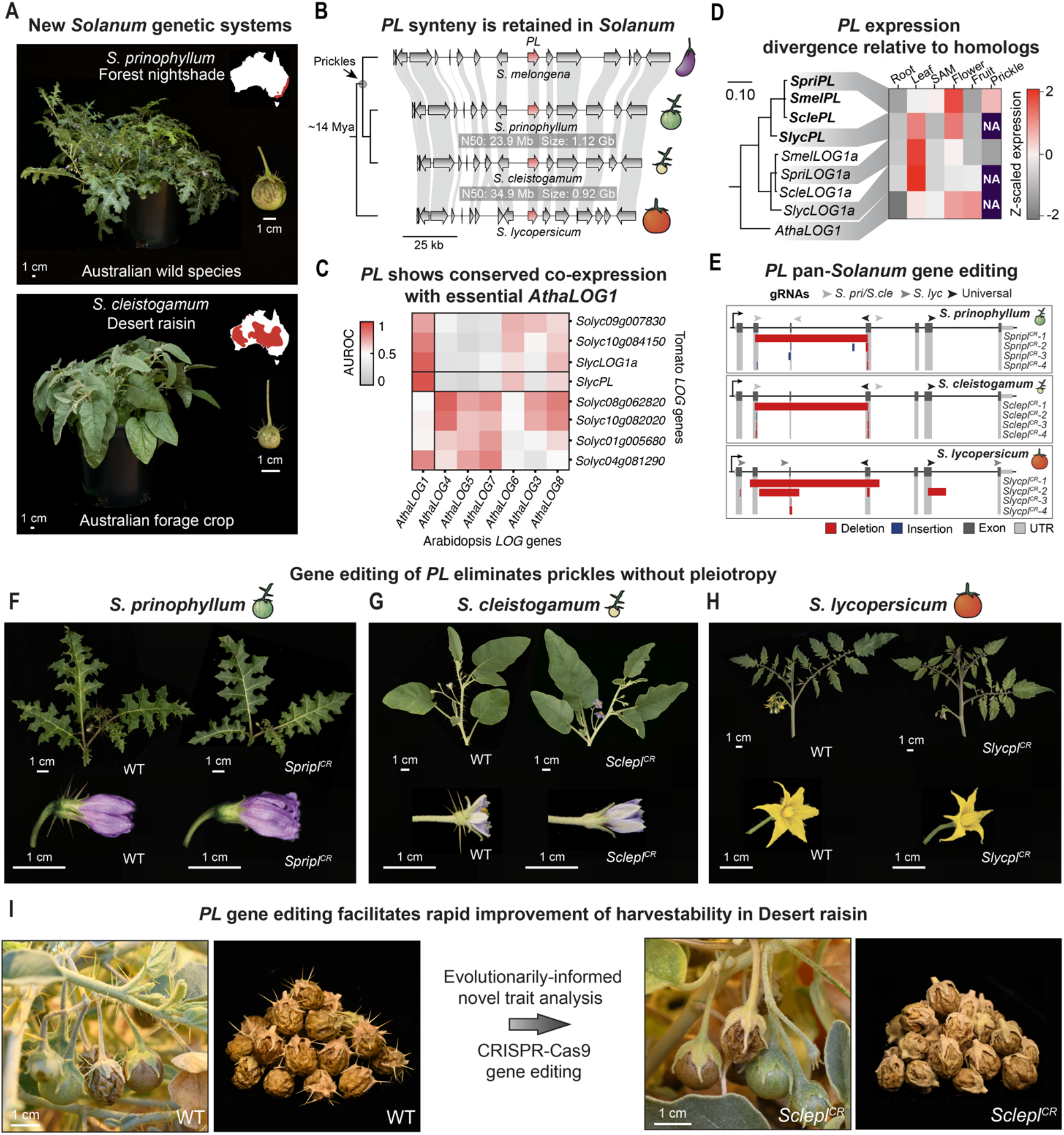
The *Solanum PL* gene was co-opted from an ancestral gene duplication event enabling non-pleiotropic editing of *PL* for crop improvement. (**A**) Whole-plant and fruit images of the prickled wild species Forest nightshade (*S. prinophyllum*, top) and its close foraged berry-producing relative Desert raisin (*S. cleistogamum*, bottom). Red-shaded region in map insets indicates approximate species ranges in Australia based on reported observations (http://www.flora.sa.gov.au/). (**B**) Genome sequencing and chromosome-scale assemblies of Forest nightshade and Desert raisin reveals that *PL* interval synteny is conserved in Brinjal eggplant and tomato (*S. lycopersicum*). Genome summary statistics are indicated. (**C**) Heatmap depicting the predictability of identifying cross-species co-expressed genes among cross-species pairs of *LOG* homologs based on their respective co-expression relationships in tomato and Arabidopsis. A higher Area Under the Receiver Operating Characteristic (AUROC) curve score indicates *LOG* homologs with increased conservation of their corresponding orthologous co-expressed genes. (**D**) Coding-sequence based maximum-likelihood phylogenetic tree of *Solanum PL* orthologs, their closely related paralog *LOG1a*, and *AthaLOG1* in comparable tissue types. Heatmap shows expression in matched tissues. (**E**) CRISPR-Cas9 targeted gene editing strategy and resulting mutant alleles generated in Forest nightshade, Desert raisin, and tomato. (**F** to **H**) Phenotypes of WT and gene edited *pl* null mutants in Forest nightshade (F), Desert raisin (G), and tomato (H). Prickles are nearly completely suppressed (Forest nightshade) and eliminated (Desert raisin) obvious pleiotropic consequences. In tomato where *PL* was not co-opted for prickle development, *Slycpl*^*CR*^ mutants resemble wild type. (**I**) Evolutionarily-informed trait analysis enables rapid and expedient removal of prickles for improved harvestability in *Solanum* crops.

Leveraging these newly developed genomic resources, we found that synteny at the *PL* locus was conserved across the *Solanum*, suggesting that *PL* was co-opted from a standing syntenic ortholog that existed at least since the divergence of tomato and the spiny *Solanums* ∼14 Mya (**Fig. 5B**). To better understand ancestral *PL* function across eudicots, we performed a meta-analysis of gene expression data from Arabidopsis (3154 samples) and tomato (5491 samples), reasoning that shared expression profiles reflect the degree of shared inter-species function (*22*). We assessed each member of the *LOG* family for its ability to predict co-expression in every other member of the *LOG* family in the other species. An area under the receiver operating characteristic (AUROC) curve statistic of 0.93 indicated that *SlycPL* in tomato is co-expressed with nearly identical genes to that of *AthaLOG1* in Arabidopsis, indicating conserved function. Likewise, three other tomato *LOG* gene family members also exhibited strongly conserved co-expression with *AthaLOG1* (**Fig. 5C**). The *LOG1* clade, to which *PL* belongs, has therefore maintained signatures of functional conservation across ∼120 My.

Tissue-specific knockdown of *AthaLOG1* in the Arabidopsis floral meristem has been shown to impair floral organ initiation, suggesting that *AthaLOG1* has critical roles in meristem maintenance, similar to the canonical developmental role for *LOGs* first reported in rice (*13, 23*). Therefore, duplication and diversification of the *LOG1* subclade in the *Solanum* may have facilitated *PL* functional co-option. To explore this hypothesis, we generated an expression atlas for prickled forest nightshade and compared it to matched-tissue gene expression data from tomato and Arabidopsis (**Fig. 5D**). In Arabidopsis, *AthaLOG1* possesses a broad expression pattern across tissues, while *Solanum PL* and *LOG1a* have evolved more tissue-biased expression patterns. Compared to its ortholog in tomato, forest nightshade *SpriPL* has evolved enriched expression in flowers and, consistent with its co-opted function, in developing prickles. Therefore, paralog diversification in the *Solanum* enabled functional co-option and redeployment of ancestral *LOG1* clade function in prickle development.

### Non-pleiotropic removal of prickles with gene editing

We reasoned that the co-option of *PL* could facilitate the engineering of agriculturally desirable loss-of-function prickleless mutants. In particular, the duplication leading to *PL* and its subsequent expression divergence from its ancestral copy would prevent undesirable pleiotropic effects on other traits. Alternatively, cryptic background modifiers in prickleless lineages may have been required to strongly suppress prickle development, and thus eliminating *PL* would leave prickles intact. To distinguish between these two possibilities, we devised a pan-genus CRISPR-Cas9 editing strategy to target *PL* in Forest nightshade, Desert raisin, and tomato, the latter of which harbors a *PL* ortholog, likely performing an ancestral function outside of prickle development. Adapting techniques previously established in tomato (*24*), we developed plant regeneration, transformation, and genome editing for Forest nightshade and Desert raisin, thereby elevating these two species into new *Solanum* genetic systems. We engineered multiple loss-of-function mutations in *PL* (*pl*^*CR*^) in all three species and compared their phenotypes (**Fig. 5E**). In both Forest nightshade and Desert raisin, *pl*^*CR*^ individuals showed strong suppression of prickle development in all tissues and organs where prickles normally develop in wild type plants, though we observed small sporadic prickles (**Figs. 5F,G and S3**). Meanwhile, in tomato, *Slycpl*^*CR*^ plants resembled the wild type, likely due to genetic redundancy with *SlycLOG1a* and possibly other *LOG* family members prior to the *PL* co-option event ∼6 Mya (**Fig. 5H**). Importantly, fruit morphology and sweetness remained unchanged (Brix sugar content ∼30% compared to ∼50% in grape raisins and ∼9% in cherry tomato) in WT and *pl*^*CR*^ Desert raisin lines, suggesting that *LOG* targeting is an effective strategy for first line improvement of harvestability in wild or partially domesticated prickled species bearing edible fruits (**Fig. 5I and Table S7**).

## Discussion

Here we showed that a repeated co-option of *LOG* family members promoted independent emergences of prickles across 150 My of plant evolution. Studies addressing convergent trait evolution at these timescales have hinted that similar and divergent genetic programs can underpin phenotypic convergence (*3, 25*). For example, the convergent evolution of echolocation in bats and cetaceans is associated with positive selection on variation in shared orthologous genes (*26*). On the other hand, different loci were reported to underlie convergent adaptation to marine habitats in mammals (*27*). The reuse of the same genetic program seen in some traits such as prickles may in part be due to their relative simplicity. Unlike composite traits (*28*), where selection has the potential to act on many different loci affecting many different organismal systems, convergent traits that arise from selection on fewer potentially relevant loci may exhibit greater genetic convergence by virtue of sheer probability. However, even traits of modest complexity, such as animal eye lenses composed of homomeric crystallins (*29*), can have many distinct genetic origins, indicating that trait complexity alone cannot not fully account for observed patterns of convergent evolution.

Genotype-phenotype convergence may also rely on developmental constraints imposed on morphological innovation, which often depends on the re-purposing of ancestral genetic mechanisms (*30, 31*). Gene co-option may allow key developmental regulators to take on new roles via non-pleiotropically partitioning gene function, particularly when standing paralog diversity exists. This has been suggested as an explanation for the repeated evolution of limbs, for example, by co-option of *Hox* genes (*32*). While cytokinin synthesis can affect essential developmental and physiological processes, functionally redundant *LOG* paralogs that arose through lineage-specific or shared ancestral duplication events (*33, 34*) may acquire specialized functions, as we found with prickles. Notably, even after their co-option in prickle development, *LOG*s may retain some functional redundancy, as engineered and natural (i.e. rice and barley) *LOG* mutants still produce sporadic small prickles (**Figs. 4A and S3**). Even partial paralog redundancy may increase the odds of phenotype-genotype convergence by allowing selection for gains and losses of prickles while avoiding developmental pleiotropy.

Lastly, as a deeply conserved plant pathway promoting cell division and differentiation (*13, 35*), cytokinin signaling has emerged as a central and repeated player in morphological innovation. For example, overexpression of a *LOG* gene is sufficient to induce the ectopic formation of shoot-borne tubers in axillary meristems in tomato (*36*), while a dominant mutation in a gene encoding a cytokinin receptor protein induces the ectopic formation of root nodules in the legume *Lotus japonicus* (*37*). However, unlike common “master” regulators that often coordinate complex programs, such as floral homeotic genes (*38*), the repeated emergence and loss of prickles reported here relies on an enzymatic gene family involved in the activation of several types of cytokinins (*39*). Hence, we hypothesize that targeted gene editing of cytokinin biosynthesis and signaling components, as demonstrated here, is likely a predictable and efficient strategy for eliminating prickles in various vascular plant lineages. This approach is particularly promising for roses, where the labor-intensive, manual removal of prickles is a common practice for most cut varieties. Continued efforts to unite genetics, genomics, and genome editing across diverse plants, as illustrated in this study, will both advance our ability to track evolutionary changes over a broad range of time scales and empower the engineering of novel phenotypes to expand our use of plant diversity in agriculture.

## Materials and Methods

### Plant growth conditions

Brinjal eggplant and wild eggplant (*Solanum insanum*) used for the mapping of *pl* were grown in a growth chamber under 25 °C/18 °C day/night temperatures and a 16/8h light/dark photoperiod, following the protocol for wild materials described in (*40*). The other eggplant species used for mapping and phenotyping were sown and germinated in 96-cell flats in a greenhouse, under natural and supplemental ∼250 μmol m^-2^ s^-1^ light from high-pressure sodium bulbs (16 h light, 8 h dark) at daytime temperatures between 26-28 °C and night-time temperatures between 18-20 °C with a relative humidity of 40-60%. At ∼4 weeks post-germination, seedlings were transplanted to 4 L pots containing high drainage PRO-MIX HP Mycorrhizae Growing Mix soil (Pro-Mix, PA, USA) supplemented with Osmocote fertilizer. Prickle phenotypes recorded for co-segregation analysis were performed in the field for the Scarlet eggplant (*S. aethiopicum*) and both the field (West Neck Field, Lloyd Harbor, NY, USA) and greenhouse for Gboma eggplant (*S. macrocarpon*). Field conditions included drip irrigation, regular soil fertilization, and manual weeding. All other *Solanum* species used in this study were grown in either the field or greenhouse, as specified (see **Table S5** for accession information).

Other plants imaged for this study were grown as follows. Rose plants were grown in the field or grown in an environmentally controlled greenhouse conditions at the Ecole Normale Supérieure of Lyon, France with 16 h light, 8 h dark day/night periods and 25 °C/19 °C day/night temperatures. Chinese date (*Z. jujuba*) images were taken of plants grown in an orchard at Los Lunas, NM, USA. Rice plants (*O. sativa*) were grown in field conditions or the greenhouse (*O. rufipogon*) and managed according to practices outlined in (*41*).

### Generation of interspecific *pl* mapping population in Brinjal eggplant (*S. melongena*)

The prickleless Brinjal eggplant *S. melongena* 8104, hereafter MEL5, and the prickled eggplant wild relative *S. insanum* SLKINS-1, hereafter INS1 (*42, 43*), were selected as founding parents to map *pl*. Advanced backcrosses (BC3) of MEL5 with introgressions from INS1 were obtained as part of a previous project (*44*).The parents, their interspecific F1 hybrid, and 90 BC3 advanced backcrosses were phenotyped for the presence of prickles at the cotyledon and 3^rd^-4^th^ true leaf development stages. Of these, 19 BC3 advanced backcrosses with prickles and 71 without prickles where genotyped with the high-throughput eggplant SPET platform (*45, 46*) to associate the presence of prickles with shared introgression intervals. After identifying the most promising genomic region, the prickled advanced backcross material with the shortest shared introgressed interval (BC3-33-3-1) was selfed to obtain a segregating BC3S1 population, which was used for fine mapping.

### Mapping of *pl* in Brinjal eggplant

A total of 622 BC3S1 seeds were germinated in growth chambers and individuals were screened for prickles two weeks post-germination. After one month, seedlings were transplanted to a greenhouse where they were phenotyped again. DNA from leaves was extracted following the SILEX protocol (*47*) and DNA yield and quality were measured spectrophotometrically using a NanoDrop™ ND-1000 (Thermo Scientific, Waltham, MA, USA). DNA integrity was checked by electrophoresis on an agarose gel (0.8%) (Condalab, Madrid, Spain) and diluted with ultra-pure water to a final concentration of 50 ng/μL. Subsequently, primers were designed for the INS1 introgression based on the genome position of target SNPs used to perform SPET in eggplant (*45, 46*) and the v3.0 eggplant genome sequence (*48*). To further delimit the ∼2.5 Mb *pl* interval and identify recombination breakpoints, we performed PCR genotyping by HRM (High Resolution Melting) genotyping (*49*). Primers used for HRM genotyping are listed in **Table S8**. PCR reactions were performed on a Roche LightCycler® 480 System (Roche Diagnostics, Rotkreuz, Switzerland) using the MasterMix qPCR No-ROX PyroTaq EvaGreen 5x for HRM (CMB-Bioline, Madrid, Spain). Two microliters of the diluted DNA sample were mixed with 0.3 μL (10 μM) of each primer, 2 μL of MaterMix qPCR qPCR No-ROX PyroTaq EvaGreen 5x for HRM and ultra-pure water until a final volume of 10 μL. The PCR reaction conditions were as follows: an initial a pre-incubation step at 95°C for 15 min followed by 55 cycles of denaturation at 95°C for 10 s, annealing at 60°C for 15 s and elongation at 68°C for 15 s. A melting curve analysis (60°C to 95°C) was performed after amplification to check specificity of the reaction. The HRM generated data were analysed using the LightCycler 480 Software release 1.5.1.62 via T_m_ values and melting curve shapes. MEL5 and INS1 parents were used as a control to detect differences between homozygote and heterozygote genotypes. The *pl* interval was thereby narrowed to a region between SNP13 and SNP14 markers, corresponding to the 105,450,271-105,543,990 bp position of chromosome 6, with a total size of 93,719 bp.

### RT-qPCR of *PL* in Brinjal eggplant (*S. melongena*)

Total RNA was isolated from the leaves of prickled and prickleless Brinjal eggplant (*S. melongena*) and prickled *S. insanum* using 700 μL of Extrazol® EM300 (Blirt DNA, Gdansk, Poland), according to the manufacturer’s specifications. Integrity was checked by 1% agarose gel electrophoresis, and purity and quantity were determined spectrophometrically using a NanoDrop™ ND-1000 (Thermo Scientific, MA, USA). Total RNA (1 μg) was treated with DNase I (RNase-free) (Thermo Scientific, MA, USA) and used as template in a FastGene Scriptase Basic cDNA Synthesis kit reaction (Nippon Genetics Europe, Düren, Germany) primed with Oligo dT. RT-qPCR was performed using a Roche LightCycler® 480 System (Roche Diagnostics, Rotkreuz, Switzerland) thermocycler using qPCR MasterMix No-ROX PyroTaq EvaGreen 5x for HRM (CMB-Bioline, Madrid, Spain). The PCR program consisted of an initial step of pre-incubation denaturation at 95°C for 15 min followed by 45 cycles of denaturation at 95°C for 15 s, annealing at 60°C for 20 s and extension at 72°C for 20 s. Melt curve analysis was performed to evaluate and confirm the specificity of the PCR reactions. An endogenous control, the eggplant housekeeping CAC gene (Clathrin adaptor complex medium subunit, SMEL_008g297560.1.01) was used as a normalization reference (*50*). Gene transcript expression level was analysed according to the relative quantitative accumulation by the 2^(-ΔΔCt)^ method (See **Table S8** for primers). A two-tailed Student’s t-test was used to determine statistical significance of inter-species differential expression.

### RT-PCR of *PL* transcripts for splice isoform analysis in Brinjal eggplant and Gboma eggplant

Approximately 0.5 cm long leaf primordia were harvested from MEL5, INS1, and a line harboring a minimal *PL* introgression from INS1 in the MEL5 background. Similarly staged tissue was also harvested from prickleless Gboma eggplant (*S. macrocarpon* PI 441914), prickled *SmacPL*^*HET*^ individuals identified in a stock of Gboma eggplant (*S. macrocarpon* SOLA112), and prickled homozygous wild-type *SmacPL* individuals in an F2 population derived from a parental cross between *S. macrocarpon* PI 441914 and *S. macrocarpon* SOLA112. All tissues were collected from greenhouse grown plants at approximately 10:00-11:00 AM and flash frozen in liquid nitrogen in 1.5 mL microfuge tubes containing a 5/32 inch (∼3.97 mm) 440 stainless steel ball bearing (BC Precision, TN, USA). Tubes containing tissue were placed in a −80 °C stainless steel tube rack and ground using an SPEX™ SamplePrep 2010 Geno/Grinder™ (Cole-Parmer, NJ, USA) for 2 min at 1440 RPM. Total RNA was extracted using TRIzol (Invitrogen, MA, USA) according to the manufacturer’s instructions for ground tissue. RNA was treated with DNase I and cleaned using an RNA Clean and Concentrator Kit (Zymo Research, CA, USA) according to the manufacturer’s instructions. Purity and concentration of the resulting total RNA was assessed using a NanoDrop One^C^ spectrophotometer (Fisher Scientific, MA, USA). Next, 1 μg of total RNA was used as input for Poly-T primed reverse transcription using the SuperScript IV VILO Master Mix (ThermoFisher, MA, USA) kit according to the manufacturer’s instructions. PCR amplification was done with 1 μL of cDNA and a primer concentration of 10 μM in a 20 μL reaction using the KOD One™ PCR Master Mix (Toyobo, Osaka, Japan). Amplification was performed using a Mastercycler X50 thermocycler (Eppendorf, Hamburg, Germany) with an initial 2 min 98 °C denaturation step, 35 cycles consisting of 10s denaturation at 98 °C, 10 s annealing at 55 °C, and 10 s extension at 68 °C, followed by a final extension step of 2 min at 68 °C. Equal volumes (5 μL) of the resulting PCR product were loaded onto a 2% biotechnology grade Agarose I (VWR International, PA, USA) gel prepared with 1X TBE buffer and 0.0025% ethidium bromide. Electrophoresis was performed in an Owl™ D3-14 electrophoresis box (Thermo Scientific, MA, USA) containing 1X TBE buffer for 45 min at 130 V delivered from an Owl™ EC 300 XL power supply (Thermo Scientific, MA, USA). The electrophoresis results were visualized under UV light using a Bio-Rad ChemiDoc™ XRS+ (Bio-Rad, CA, USA) imaging platform and ImageLab™ (Bio-Rad, CA, USA) software. Relevant primer sequences are listed in **Table S8**.

To determine the identity and relative abundance of transcript isoforms across the different genotypes, PCR products were cloned using a StrataClone Blunt PCR Cloning Kit (Agilent, CA, USA) according to the manufacturer’s instructions. PCR was conducted on single colonies swabbed into 10 μL sterile water and a 1 μL aliquot of the suspended culture was used as a PCR reaction template using 10 μM M13F/R primers with KOD One™ PCR Master Mix. PCR amplification was performed with a 2 min 98 °C denaturation step, 35 cycles consisting of 10 s denaturation at 98 °C, 10 s annealing at 55 °C, and 10 s extension at 68 °C and followed by a final extension step of 2 min at 68 °C using a Mastercycler X50 thermocycler. PCR products were purified using 1.8X volume Ampure XP beads according to the manufacturer’s instructions and submitted for Sanger sequencing (Azenta Genewiz, NJ, USA).

### Co-segregation and mapping of *pl* in Scarlet eggplant (*S. aethiopicum*) and Gboma eggplant (*S. macrocarpon*)

The *SaetPL* allele co-segregation analysis in Scarlet eggplant was performed on an F2 population derived from an intraspecific cross between the prickled accession SC102 and the prickleless accession PI 424860. Field grown plants were phenotyped for the presence/absence of prickles and young leaf tissue was collected and DNA extracted using the cetrimonium bromide (CTAB) method (*51*). Primers (see **Table S8**) were designed to amplify the exonic region of *PL* harboring the mutant *Saetpl* deletion allele identified in the genome of the prickleless parental accession. PCR reactions were prepared using 1 μL of DNA and 10 μM primers in a 10 μL reaction volume with KOD One™ PCR Master Mix (Toyobo, Osaka, Japan). Amplification was performed on a Mastercycler X50 thermocycler (Eppendorf, Hamburg, Germany) with a 2 min 98 °C denaturation step, 35 cycles consisting of 10 s denaturation at 98 °C, 10 s annealing at 55 °C, and 10 s extension at 68 °C, followed by a final extension step of 2 min at 68 °C. PCR products were purified using a 1.8X volume of AMPure XP beads (Beckman Coulter, CA, USA) and submitted for Sanger sequenced (Azenta Genewiz, NJ, USA) for *SaetPL* genotyping. Genotypes were then associated with prickle phenotypes. Chi-squared goodness of fit tests were performed with GraphPad.

For *SmacPL* in Gboma eggplant the same initial co-segregation analysis was performed as was done for Scarlet eggplant, above (see **Table S8** for primers). Multiple F2 populations derived from an intraspecific cross between prickleless Gboma eggplant (*S. macrocarpon* PI 441914) and prickled Gboma eggplant (*S. macrocarpon* SOLA112) were generated. QTL-Seq was performed on a single F2 population made up of the progeny of a single prickled F1 parent. From a population of 132 individuals, the DNA from a random selection of 25 prickleless and 31 prickled F2 individuals, along with 8 prickleless parental accession individuals (*S. macrocarpon* PI 441914) was extracted from young leaf tissue using a DNeasy Plant Pro Kit (Qiagen, Hilden, Germany) according to the manufacturer’s instructions for high-polysaccharide content plant tissue. Tissue used for extraction was ground using an SPEX™ SamplePrep 2010 Geno/Grinder™ (Cole-Parmer, NJ, USA) for 2 min at 1440 RPM. Sample DNA (1 μL assay volume) concentrations were quantified with Qubit 1X dsDNA HS buffer (ThermoFisher, MA, USA) on a Qubit 4 fluorometer (ThermoFisher, MA, USA) according to the manufacturer’s instructions. Separate pools were made for the parental sample, the prickled F2 individuals, and the prickleless F2 individuals, with an equivalent mass (3 μg) of DNA pooled from each individual. Next, the DNA pools were purified using a 1.8X volume of AMPure XP beads (Beckman Coulter, CA, USA) and the DNA concentration and purity assayed by Qubit and a NanoDrop One^C^ spectrophotometer (Fisher Scientific, MA, USA), respectively.

Paired-end sequencing libraries for QTL-Seq analysis were prepared using >1 μg of pooled DNA with a KAPA HyperPrep PCR-free kit (Roche, Basel, Switzerland) according to the manufacturer’s instructions. Indexed libraries were pooled for sequencing on a NextSeq 2000 P3 chip (Illumina, CA, USA). Mapping was performed using using the end-to-end pipeline implemented in the QTL-Seq software package (v2.2.4, https://github.com/YuSugihara/QTL-seq) (*52*) with reads aligned against the *S. macrocapron* (Smac3, PI 441914) genome assembly. After QTL-Seq analysis revealed a second interval, *pl2*, an additional co-segregation analysis was performed across multiple F2 populations, genotyping *SmacPL* and the *pl2* locus on chromosome 4 identified by QTL-Seq. For the *pl2* interval, an A-to-T SNP at position Chr4:20,401,670, which modifies a DraI restriction site, was used to as the basis for a cleaved amplified polymorphic sequences (CAPS) genotyping assay. Following PCR amplification using the same conditions described for the *SaetPL* genotyping above (see **Table S8** for primer details), 5 μL of PCR product was added to a 10 μL reaction containing 0.2 μL DraI (New England BioLabs, MA, USA) and 1 μL rCutSmart™ Buffer (New England BioLabs, MA, USA) and incubated for 2-4 hours at 37 °C. The reactions were then loaded onto a 1% agarose gel and electrophoresed in an Owl™ D3-14 electrophoresis box (Thermo Scientific, MA, USA) containing 1X TBE buffer for 30 min at 180 V delivered from an Owl™ EC 300 XL power supply (Thermo Scientific, MA, USA). The electrophoresis results were visualized under UV light using a Bio-Rad ChemiDoc™ XRS+ (Bio-Rad, CA, USA) imaging platform and ImageLab™ (Bio-Rad, CA, USA) software. Genotypes assigned based on visualized PCR product cleavage patterns and associated with prickle phenotypes. Chi-squared goodness of fit tests were performed with GraphPad.

### Genome assembly and scaffolding

Reference quality genome assemblies for *S. aethiopicum, S. macrocarpon, S. prinophyllum*, and *S. cleistogamum* (see **Table S5** for accession information) were generated using a combination of long-read sequencing (Pacific Biosciences, CA, USA) for contigging and optical mapping (Bionano Genomics, CA, USA) for scaffolding. High-molecular weight DNA used for sequencing was extracted from greenhouse grown 4 week-old seedlings germinated in 96-cell flats and dark treated for 48 hrs prior to flash freezing. High molecular weight DNA was extracted according to the method described in (*53*). Two PacBio Sequel IIe flow cells (Pacific Biosciences, CA, USA) were used for the sequencing of each sample (average read N50 = 11,221 bp, average coverage = 53X, average read QV = 83.28). Prior to assembly, we counted k-mers from raw reads with KMC3 (version 3.2.1) and estimated genome size, sequencing coverage, and heterozygosity with GenomeScope2.0 (*54*). Sequencing reads from each sample were assembled with hifiasm (*55*) exact parameters and software version varied between samples based on the level of estimated heterozygosity and are reported in **Table S3**. Post assembly, the draft contigs were screened for possible microbial contamination as described in (*53*).

Optical mapping (Bionano Genomics, CA, USA) was performed for each sample to facilitate scaffolding. Scaffolding with optical maps was performed using the Bionano solve Hybrid Scaffold pipeline with default parameters (https://bionanogenomics.com/support/software-downloads/). Hybrid scaffold N50s ranged from 75,028,200 bp to 107,486,353 bp (see **Table S3** for more detail including Bionano molecules per sample). High-throughput chromosome conformation capture (Hi-C) from Arima Genomics, CA, USA was performed for one sample, *S. prinophyllum*, to finalize scaffolding. 311,616,288 reads were integrated with the Juicer (v0.7.17-r1198-dirty) pipeline. Next, misjoins and chromosomal boundaries were manually curated in the Juicebox (v1.11.08) application. Chromosomes were named based on sequence homology, determined with RagTag (*56*) scaffold (v2.1.0, default parameters), with the phylogenetically-closest finished genome (see **Table S3** for details). Finally, small contigs (< 50,000 bp) with > 95% of the sequence mapping to a named chromosome were removed. Additionally, small contigs (< 100,000 bp) with > 80% of the sequence mapping to a named chromosome that contained one or more duplicated BUSCO genes, but no single BUSCO genes, were also removed using a python script. Using merqury (*57*) with the HiFi data, the final consensus quality of the assemblies was estimated as QV=51.1333 on average and a completeness of 99.2741% on average.

### Genome annotation

The gene annotation pipeline involved several crucial steps. Initially, the quality of raw RNA-Seq reads underwent assessment using FastQC v0.11.9. Subsequently, reference-based transcripts were generated using STAR v2.7.5c (*58*) and Stringtie2 v2.1.2 (*59*) workflows. To refine the data, invalid splice junctions from the STAR aligner were filtered out utilizing Portcullis v1.2.0 (*60*). Orthologs with coverage above 50% and 75% identity were lifted from Heinz v4.0 (*61*) and Eggplant v4.1 (*62*) via Liftoff v1.6.3 (*63*) using parameters --copies,--exclude_partial and employing both Gmap version 2020-10-14 (*64*) and Minimap2 v2.17-r941 (*65*) aligners. In addition, protein evidence from several published Solanaceae genomes (*61, 62, 66*), and the UniProt/SwissProt database were utilized to support gene annotation. Structural gene annotations were generated through the Mikado v2.0rc2 (*67*) framework, leveraging evidence from the Daijin pipeline (*68*). Additionally, microsynteny and orthology to Heinz v4.0 and Eggplant v4.0 were assessed using Microsynteny and Orthofinder v2.5.2 (*69*). Correction of gene models with inframe stop codons utilized Miniprot2 (*70*) protein alignments from Heinz v4.0 and Eggplant v4.1. Furthermore, gene models lacking start or stop codons were adjusted by placing them within 300 base pairs of the nearest codon location using a custom python script.

For functional annotation, ENTAP v0.10.8 (*71*) integrated data from diverse databases such as PLAZA dicots (5.0) (*72*), Uniprot/Swissprot (*73*), TREMBL, RefSeq, Solanaceae proteins, and InterProScan5 (*74*) with Pfam, TIGRFAM, Gene Ontology, and TRAPID (*75*) annotations. Finally, the annotated data underwent a series of filtering steps, excluding proteins shorter than 20 amino acids, those exceeding three times the length of functional orthologs, proteins lacking assigned orthologs that have unknown function, and transposable element (TE) genes, which were removed using the TEsorter (*76*) pipeline.

The completeness of the gene models was determined by assessing single-copy orthologs through BUSCO5 (*77*) in protein mode, comparing against the solanales_odb10 database. Additionally, the presence or absence of a curated set of 180 candidate genes known to be crucial in QTL studies was examined. Genome annotation summary statistics are presented in **Table S4**.

DNA extraction, *PL* PCR amplification, and sequencing of *Solanum* species

For herbarium samples, tissue (∼ 1-2 cm^2^) was excised with a scalpel from mounted herbarium specimens held in the Steere Herbarium Collection at the New York Botantical Garden, New York, USA (see **Table S5** for voucher information). Silica gel-preserved tissues were obtained from the Royal Botanic Garden Edinburgh. Tissue was obtained from species documented to be prickleless and samples were inspected for the absence of prickles. Likewise, tissue was also obtained from closely-related prickled sister species for sequence comparison. In total, 8 putative loss-of-function coding or splice-site mutations were identified in herbarium samples across 7 species. Two distinct mutations were identified in separate *S. schimperianum* specimens. For technical reasons, it was not possible to survey the complete exonic space of all genes - primer binding site divergence and low DNA sample quality precluded complete survery exonic survey in most cases. Among prickleless herbarium specimens in which a complete exonic survery was possible, 7 lacked an obvious putative loss-of-function coding or splice-site mutation. Among the remaining prickleless herbarium material with incomplete exonic sequence coverage, 22 specimens had no detectable putative loss-of-function coding or splice-site mutations. However, this does not preclude the possibility that *PL* mutations exist in these samples. Such mutations could lie in unsampled regions of the gene body or in non-coding regions. In addition, other loci (i.e. *pl2* in *S. macrocarpon*) may also contribute to unexplained prickle losses in *Solanum*. Meanwhile, across 30 sampled prickled sister taxa, none of the reported loss-of-function *pl* alleles were found and no other putative loss-of-function coding or splice-site mutations identified.

To extract DNA from herbarium samples, ∼25 mg of tissue was added to a 2 mL microfuge tube containing a 5/32 inch (∼3.97 mm) 440 stainless steel ball bearing (BC Precision, TN, USA) and ground for 2 min at 1440 RPM in an SPEX™ SamplePrep 2010 Geno/Grinder™ (Cole-Parmer, NJ, USA). Before beginning the extraction the bench area and pipettes were cleaned with a 10% bleach solution to reduce the risk of environmental DNA contamination. DNA extraction was performed using a modified version of the CTAB/STE-based extraction methods described in (*78*) and (*79*). First, 1 mL of freshly-prepared STE (0.25 M sucrose, 0.03 M Tris pH 8, 0.05 M ethylenediaminetetraacetic acid [EDTA]) was added to the ground tissue and vortexed for 5 s. The sample was then centrifuged at 2000 x g for 10 min and the wash and centrifugation in STE buffer was repeated once. Next, 1 mL 2X CTAB and 4 μL of 1 M dithiothreitol (DTT) were added to the precipitated tissue and incubated with gentle rotating agitation in a hybridization oven for 1 hr at 65 °C. Following incubation, organic extraction was performed by addition of 750 μL of 24:1 chloroform/isoamyl alcohol and the samples were inverted 10 times. Phase separation was achieved by centrifugation for 10 min at 16,200 x g and ∼900 μL of the aqueous phase was transferred to a new tube. To precipitate the DNA, 3 μL of 15 mg/mL GlycoBlue™ coprecipitant (ThermoFisher Scientific, MA, USA) was added and well-mixed followed by 600 μL of chilled −20 °C isopropanol. DNA was allowed to precipitate at −20 °C for 1 hr. After precipitation, the samples were centrifuged at 20,000 x g for 30 min in a centrifuge chilled to 4 °C. The resulting DNA pellet was washed once with 70% ethanol and re-suspended in 50-100 μL 0.1X TE pH 8 buffer.

Because DNA form archived dried plant tissues is often fragmented (*78*), an exon screening based approach was taken to identify mutations in the CDS or immediately flanking splice acceptor/donor sites. Primers were designed for intronic regions ∼20-30 bp surrounding exons based on the sequence of *PL* from closely-related species for which whole-gene amplification of *PL* was possible due to the availability of intact DNA isolated from fresh, living tissue (see **Table S8** for primers). PCR amplification was performed with 1-2 μL of DNA, primers at 10 μM, in a 10 μL reaction using the KOD One™ PCR Master Mix (Toyobo, Osaka, Japan). Amplification was performed using on a Mastercycler X50 thermocycler (Eppendorf, Hamburg, Germany) with a 2 min 98 °C denaturation step, 35-40 cycles consisting of 10 s denaturation at 98 °C, 10 s annealing at 48-62 °C, and 10 s extension at 68 °C, followed by a final extension step of 2 min at 68 °C. PCR products were purified using 1.8X volume Ampure XP beads (Beckman Coulter, MA, USA) according to the manufacturer’s instructions. For high-yielding PCR reactions, the PCR products were directly Sanger sequenced (Azenta Genewiz, NJ, USA). For low-yielding PCR reactions, the PCR products were blunt cloned and sequenced as done for the PL transcript splicing isoform analysis described above. Sequences were then aligned (Geneious alignment) to either Brinjal eggplant *PL* gene body and screened for mutations using Geneious version 2022-07-07 software (see **Table S6**).

For the genotyping of *PL* in prickleless species for which fresh tissue was available, a CTAB-based method was used to obtain DNA (*51*). Primers (see **Table S8**) anchored in the first and final exon of *PL* were used to PCR amplify the entire gene body. Touchdown PCR amplification was performed using on a Mastercycler X50 thermocycler (Eppendorf, Hamburg, Germany) with an initial 2 min 98 °C denaturation step, 16 cycles consisting of an initial 10 s annealing at 68 °C thereafter descending 1 °C/cycle, a 1 min extension at 68 °C, and a 10 s 98 °C denaturation. Next, the reaction proceeded through 14-19 cycles of annealing for 10 s at 60 °C, extension for 1 min at 68 °C and 10 s denaturation at 98 °C followed by a final extension of 2 min at 68°C. PCR products were purified using 1.8X volume Ampure XP beads (Beckman Coulter, MA, USA) according to the manufacturer’s instructions. PCR products were then blunt cloned, as described above, and whole-plasmid sequenced (Plasmidsaurus, OR, USA) to detect variants across the *PL* gene body. In all 4 such cases, candidate *PL* mutations were identified. See **Table S6** for identified exonic *PL* mutations in prickleless species and the sequences of aligned prickled sister taxa).

### Analysis of published mapping and RNA-Seq data from rose

The major QTL for prickle formation on rose stems was previously mapped on chromosome 3 using a mapping population derived from a cross between of *R. chinensis* ‘Old Blush’ (OB) and *R. wichurana* Basye’s Thornless (BT) (*8, 80*). All genes in the Prickles QTL interval were *de novo* annotated in the genome of *R. wichurana* BT using the gene prediction tool AUGUSTUS (*81*). Predicted annotations were refined by mapping the OB protein to the BT genomic regions with Exonerate (*82*). Synteny is based on the match found by Exonerate. Previously published RNA sequencing data were used to examine expression of the LOG homolog contained within the rose mapping interval. Reads were trimmed using trimmomatic v0.39 (*83*) and then mapped to the Rosa chinensis genome using STAR v2.7.5c (*58*). Read pileups at 1 bp resolution were visualized by converting resulting bam files to bigwig format.

### Identification of LOG mutations in Chinese date (*Ziziphus jujuba*) and giant spider-flower (*Tarenaya hassleriana*)

Published genomes of Chinese date (*Z. jujuba*) were screened for mutations in LOG homologs. Candidate exonic loss-of-function mutations were identified in the cultivated spine-suppressed *Z. jujuba* cultivar Dongzao (GenBank assembly: GCA_000826755.1) and *Z. jujuba* cultivar Junzao (GenBank assembly: GCA_001835785.2) in a *LOG* homolog corresponding to LOC107411352. Mutations were not detected in this gene in the spiny wild relative *Z. jujuba* var. spinosa (GenBank assembly: GCA_020796205.1). For giant spider-flower a mutation was identified in a *LOG* homolog (LOC104826905) in the reference genome (GenBank assembly: GCA_000463585.1).

### Phylogenetic analysis of angiosperm *LOG* homolog

*LOG* genes were identified across plant genomes using Orthofinder (*69*). for the chosen eleven species, the protein sequences of ninety-nine proteins were aligned using MAFFT v7.402 (*84*). The best scoring maximum likelihood tree was inferred using default parameters, with 1000 bootstrap replicates in RAxML v8.2.12 (*85*) via CIPRES (*86*). The tree was visualized in R using the ggtree package (*87*). The AtLOG_TypeIIIb (AT1G50575.1) protein sequence was used as an outgroup. Branch support was provided by 1000 replicate bootstrapping. The *LOG* sub-clades were designated after the naming convention used in (*34*).

### Forest nightshade (*S. prinophyllum*) expression atlas

Washed and dried roots, leaves, flowers, fruits (with seeds removed), and developing prickles (∼0.25-0.50 cm in length) were collected from 2 month-old Forest nightshade (*S. prinophyllum*) plants. All tissues were collected in 3-4 replicates. Total RNA was extracted using Quick-RNA MicroPrep Kit (Zymo Research) and treated with DNAse I (Zymo Research) according to the manufacturer’s instructions. RNA concentration and quality were analyzed using Thermo Scientific™ NanoDrop™ One^C^ Spectrophotometer. Libraries for RNA-sequencing were prepared by KAPA mRNA HyperPrep Kit (Roche, Basel, Switzerland). Paired-end 100-base sequencing was conducted on the NextSeq 2000 P3 sequencing platform (Illumina, CA, USA). Reads were trimmed using trimmomatic v0.39 (*83*) and then mapped to the Spri1 genome using STAR v2.7.5c and expression computed in transcripts per million (TPM). Expression data from the Forest nightshades in vegetative meristem generated in a previous study was processed in the same fashion (*88*). Gene expression data from ATHENA (https://athena.proteomics.wzw.tum.de/master_arabidopsisshiny/) for the *Arabidopsis LOG1* ortholog in equivalent tissues was downloaded and data for LOG homologs in tomato was downloaded from https://www.solpangenomics.com. Expressed genes were clustered phylogenetically based on alignment of their CDS sequences using the Clustal Omega function in Geneious (version 2022-07-07). A maximum likelihood tree using the Tamura-Nei model (*89*) was used for tree building.

### Generation of CRISPR-Cas9-induced mutants and phenotyping

CRISPR Guide RNAs to target *PL* across Solanum species were designed using Geneious. The Golden Gate cloning approach as described in (*24*) was used to create multiplexed gRNA constructs. Plant regeneration and *Agrobacterium tumefaciens*-mediated transformation of *S. prinophyllum* and tomato were performed according to (*90*). The same methods were also used for *S. cleistogamum* with two modifications. First, for plant regeneration, the medium was supplemented with 0.5 mg/L zeatin instead of 2 mg/L and for the selection medium, 75 mg/L kanamycin was used instead of 200 mg/L. Seed germination time in culture can vary between species and batches of harvested seeds. Typically, *S. prinophyllum* germination took 8 – 10 days and *S. cleistogamum* behaved similarly to tomato, germinating in 6 - 8 days.

First generation transgenic (T0) plants were screened for prickle suppression phenotypes relative to non-transgenic regenerated control plants. DNA was extracted from T0 individuals using the CTAB method (*51*) and primers (see **Table S8**) were used to amplify the entire *PL* gene body, which was cloned, and whole-plasmid sequenced as was done for the pan-Solanum screening for mutations in the *PL* gene body. For each T0 individual, 5-8 clones were sequenced to verify the association of the prickleless phenotype and null multiallelic or monoallelic T0 phenotype. For *S. prinophyllum* and *S. lycopersicum*, CRISPR alleles advanced to the T1 generation showed similar phenotypes to the T0.

### Scanning Electron Microscopy

For prickle imaging in barley (*Hordeum vulgare*), freshly isolated awn samples were fixed with 2% glutaraldehyde in 50 mM phosphate buffer pH 7.0 for 16h at 8°C. After two 5 min washes with distilled water, samples were dehydrated in an ascending ethanol series of 30%, 50% 70%, 90%, 100%, and a second 100% ethanol wash with each step lasting 10 min after which they were critical point dried in a Quorum K850 critical point dryer (Quorum Technologies Ltd., https://www.quorumtech.com). Dehydrated samples were placed onto carbon adhesive discs, gold coated in an Edwards S150B sputter coater (Edwards High Vacuum Inc., Burgess Hill, UK) and examined in a Zeiss Gemini300 scanning electron microscope (Carl Zeiss Microscopy GmbH, Jena, Germany) at 5 kV acceleration voltage. For scanning electron microscopy of rose (*Rosa*), *Solanum*, and rice (*Oryza sativa* and *O. rufipogon*) prickles, fresh tissue was adhered to stubs using non-conducting adhesive tape and imaged using a NanoScope JCM-7000 Benchtop SEM (JEOL, Tokyo, Japan).

### Brix assay to measure fruit sugar content

From 6 Desert raisin (*S. cleistogamum*) plants, 12 fully ripened and dried fruits were randomly selected to determine their soluble sugar content (Brix). Fruits were briefly rehydrated by soaking in DI water for 60 minutes. Following rehydration, the fruit were patted dry and placed into a 50 mL tube. Rehydrated fruit were then crushed using a spatula. The slurry was centrifuged at 15,000 rpm for 5 min. The supernatant was drawn from the samples and diluted 1:30 to obtain a working volume of 300 μL. The Brix value (%) was quantified from three technical replicates with an ATAGO Palette digital Brix refractometer (ATAGO, Tokyo, Japan). See **Table S7** for data.

### Cross-species *LOG* family co-expression conservation analysis

To calculate the co-expression conservation between tomato genes and *Arabidopsis thaliana* genes, gene family orthology information was downloaded from OrthoDB V11(*91*). Using one-to-one focal gene pairs between the two species, the degree to which co-expression is conserved between all *LOG* genes in each species was calculated (*92*). Briefly, a robust gold standard coexpression network for each species was sourced (*22*), which consists of coexpression networks built from hundreds of samples across dozens of experiments. For each Arabidopsis *LOG* gene, the top 10 genes it was coexpressed with were identified and used to predict the coexpression partners of all *LOG* genes in tomato. The degree to which the top coexpressed genes in *Arabidopsis* predicted the top coexpressed genes for a given *LOG* homolog in tomato yielded the co-expression conservation score, which is defined as the area under the receiver operatoring characterstic (AUROC) curve. This was then repeated in the other direction, taking the top 10 coexpression partners of each tomato *LOG* gene and predicting the coexpression partners of all Arabidopsis *LOG* genes. The two resulting AUROC statistic values are averaged, resulting in the final co-expression conservation scores.

## Acknowledgements

We thank members of the Lippman laboratory and critical friend M. Bartlett for discussions and feedback on the manuscript. We acknowledge B. Semen and R. Santos from the Lippman lab for technical support, T. Mulligan, K. Schlecht, S. Qiao for assistance with plant care, S. Goodwin, S. Eskipehlivan, and D. McCombie for next-generation sequencing services, M. Frank and N. Lippman for photographs of *V. amazonica*, J. Emerson for his photograph of *S. pyrocanthos*, students in the Frary laboratory at Mt. Holyoke College for validation of genome edited plant phenotypes, and H. Golan for his work developing solpangenomics.com. We thank N. Tarnowsky, M. Pace, and the New York Botanical Garden for facilitating tissue sampling at the Steere Herbarium and reproduction of digitized collections held in the C.V. Starr Virtual Herbarium. We acknowledge the USF Institute for Systemic Botany for use of the digitized herbarium image of *T. hassleriana* and the use of public domain photos from R. Hannawacker of *S. greggii* and Zayda C. of *D. stramonium*. We express our gratitude to the Peruvian government for permission to collect and sequence indigenous *Solanum* species (Ministerio de Agricultura, Dirección General Forestal y de Fauna Silvestre collection permits 084-2012-AG-DGFFSDGEFFS and 096-2017-SERFOR/DGGSPFFS, and genetic resource permit 008-2014-MINAGRI-DGFFS/DGEFFS). We are also grateful to our colleagues L.L. Giacomin and J.R. Stehmann for providing access to their *Solanum* collections and the Brazilian government for granting collecting and export permits (Cadastro A456CF7) for the RPPN Santuário do Caraça and Alto da Serra de Paranapiacaba. Lastly, we gratefully acknowledge the First Nations people of Australia for their care of the land and biodiversity where *S. cleistogamum* and *S. prinophyllum* were originally collected.

## Author contributions

**Conceptualization:** JWS, ZBL; **Formal analysis:** MJP, JG; **Funding acquisition:** JWS, PG, AA, GV, MB1, MP, EG, RH, NS, MB2, SK, TS, JG, JVE, MCS, JP, ZBL; **Investigation:** JWS, DA, PG, KMJ, JH, AA, GV, MP, SR, MB1, IG, AH, HS, BF, GMR, YG, KS, MJP, EG, RH, SK, TS, JG, JVE, MCS, JP, SV, ZBL; **Methodology:** JWS, DA, PG, KMJ, AA, GV, MP, SR, IG, AH, BF, YG, KS, MJP, SK, TS, JG, JVE, MCS, JP, SV, ZBL; **Project administration:** JWS, MCS, JP, ZBL; **Resources:** JWS, DA, PG, KMJ, JH, AA, GV, MP, SR, MB1, IG, AH, HS, BF, GMR, YG, KS, MJP, EG, RH, TDH, GCE, AG, TR, NS, SY, CB, MB2, AF, SK, TS, JG, JVE, MCS, YE, JP, SV, ZBL; **Software:** KMJ, SR, MJP, JG, MCS; Supervision: JP, SV, ZBL; **Validation:** JH, TDH, GCE, TR, NS, CB, MB2; **Visualization:** JWS, DA, PG, MJP, JG, TS, JP, SV, ZBL; **Writing – original draft:** JWS, ZBL; **Writing – review & editing:** JWS, AF, YE, ZBL.

## Funding

National Science Foundation Postdoctoral Fellowship in Biology [IOS-2305651) (to **JWS**)]; Postdoctoral fellowship (RYC2021-031999-I) funded by MCIN/AEI /10.13039/501100011033 and by “European Union NextGenerationEU/PRTR” (to **PG**); Predoctoral fellowship (FPU18/01742) from Spanish Ministerio de Ciencia, Innovación y Universidades (to **AA**); Predoctoral fellowship (PRE2019-086256) funded by MCIN/AEI/10.13039/501100011033 and by “ESF investing in your future” (to **GV**); Biologie et amélioration des Plantes departments of the French National Institute for Agriculture, Food, and Environment [INRAE (to **MB1, MB2**)]; The William Randolph Hearst Scholarship from the Cold Spring Harbor School of Biological Sciences (to **MJP**); Fonds de recherche du Québec en Nature et Technologies Postdoctoral Fellowship (to **EG**); The Sibbald Trust Research Fellowship (to **RH**); Grant of the German Research Association DFG [STE 1120/17-1 (to **NS**)]; NSF Planetary Biodiversity Initiative grant “PBI Solanum: a worldwide initiative” [DEB-0316614 (to **SK**)]; National Geographic Society Northern Europe Award GEFNE49-12 (to **TS**); National Institutes of Health grant R01MH113005 (to **JG**); Grant CIPROM/2021/020 funded by Conselleria d’Innovació, Universitats, Ciència i Societat Digital of the Generalitat Valenciana (to **JP**); Grant PID2021-128148OB-I00 funded by MCIN/AEI/10.13039/501100011033/ and “ERDF A way of making Europe” (to **JP**); Grant PDC2022-133513-I00 funded by MCIN/AEI/10.13039/501100011033/ and by “European Union NextGenerationEU/PRTR” (to **JP**); National Science Foundation Plant Genome Research Program grant IOS-2216612 (to **AF, JG, JVE, MCS, ZBL**); The Howard Hughes Medical Institute (to **ZBL**).

---

**Fig. S1.**
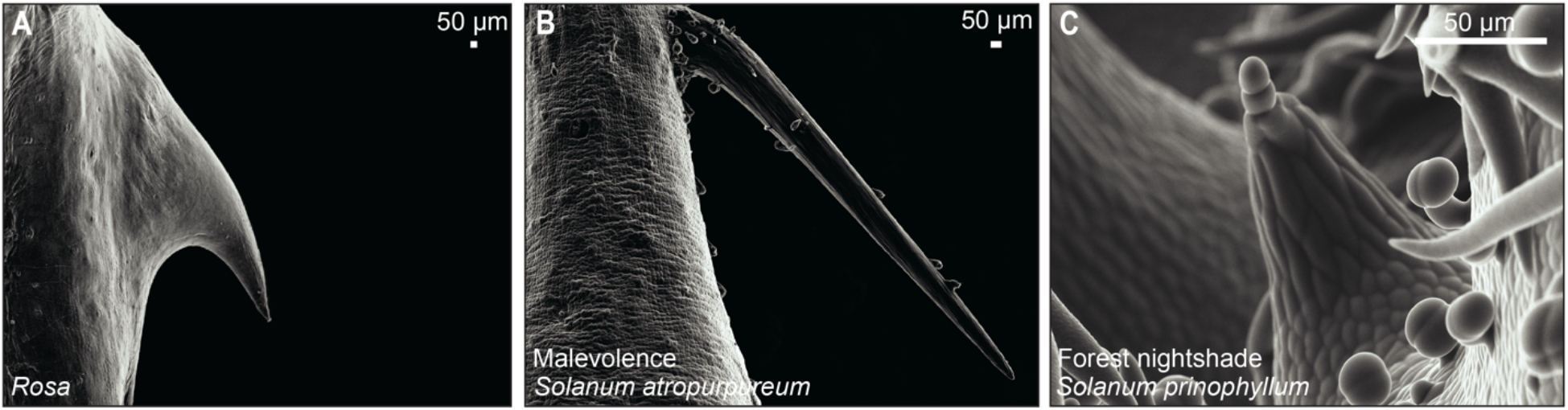
Prickles are morphologically similar. (**A-C**) Scanning electron micrographs (SEMs) of prickles in rose (*Rosa* spp.) (A), Malevolence (*Solanum atropurpureum*) (B), and at an early stage of development in Forest nightshade (*Solanum prinophyllum*) (C).

**Fig. S2.**
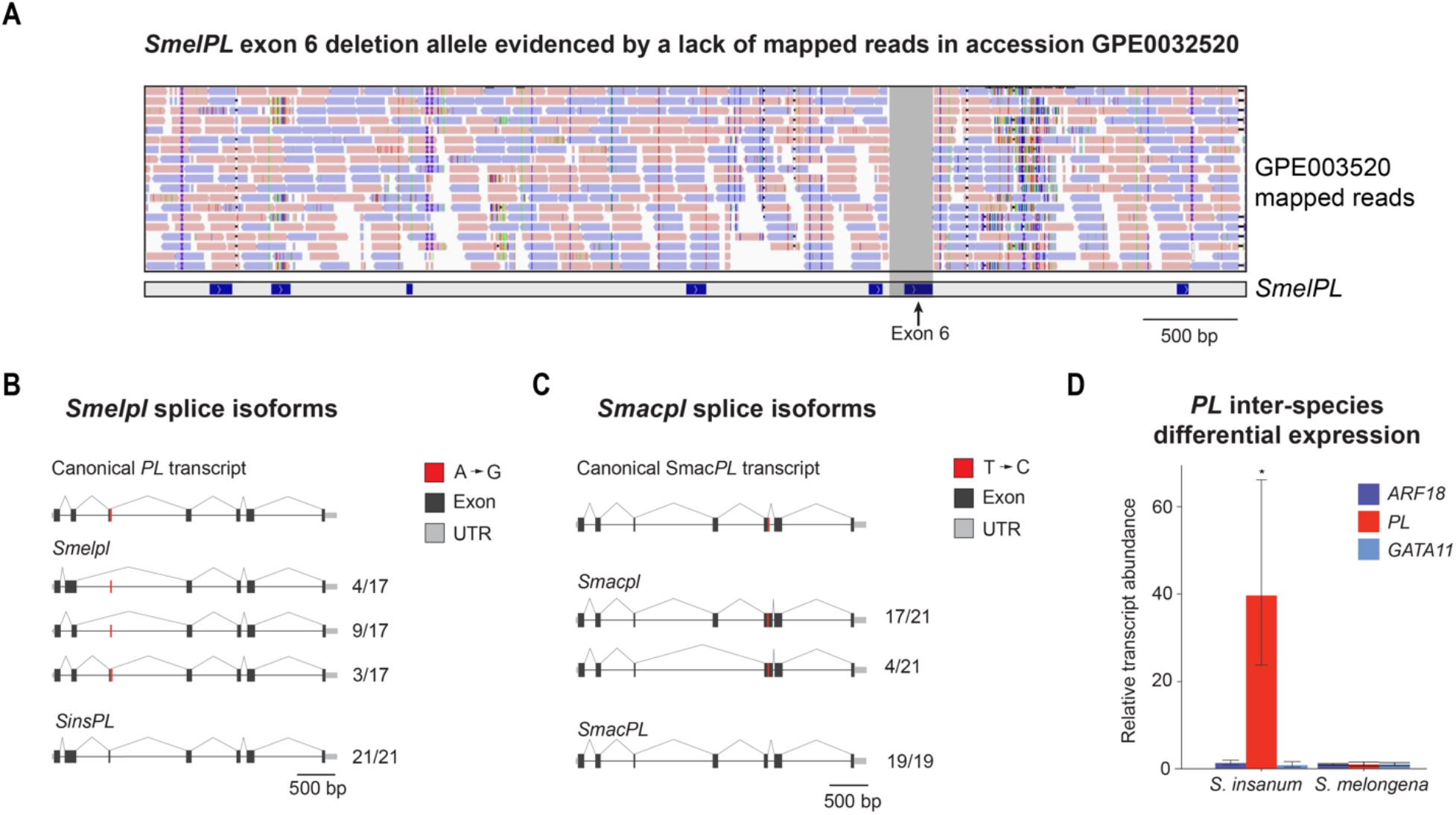
Characterization of *PL* alleles in Brinjal (*S. melongena*) and Gboma eggplant (*S. macrocarpon*). (**A**) A second *Smelpl* allele in the re-sequenced accession GPE003520 removed exon 6 as evidenced by a lack of whole-genome sequencing reads mapped to this region. Reads mapping to the forward strand are in red reads mapping to the reverse strand are in blue. (**B, C**) Gene body diagrams illustrate the canonical and *PL* isoforms for common eggplant (A) and Gboma eggplant (B). Below are detected *PL* isoforms from the leaf tissue of plants homozygous for the indicated alleles. Fractions represent the proportion of isoforms of each type detected. (**D**) RT-qPCR analysis of *PL* expression in the leaves of wild eggplant (*S. insanum*) and common eggplant. The expression of two transcription factors nearby the *pl* mapping interval are also shown. Error bars reflect the standard deviation about the mean, Student’s two-tailed t-test, * *p* < 0.05, *n* ≥ 3.

**Fig. S3.**
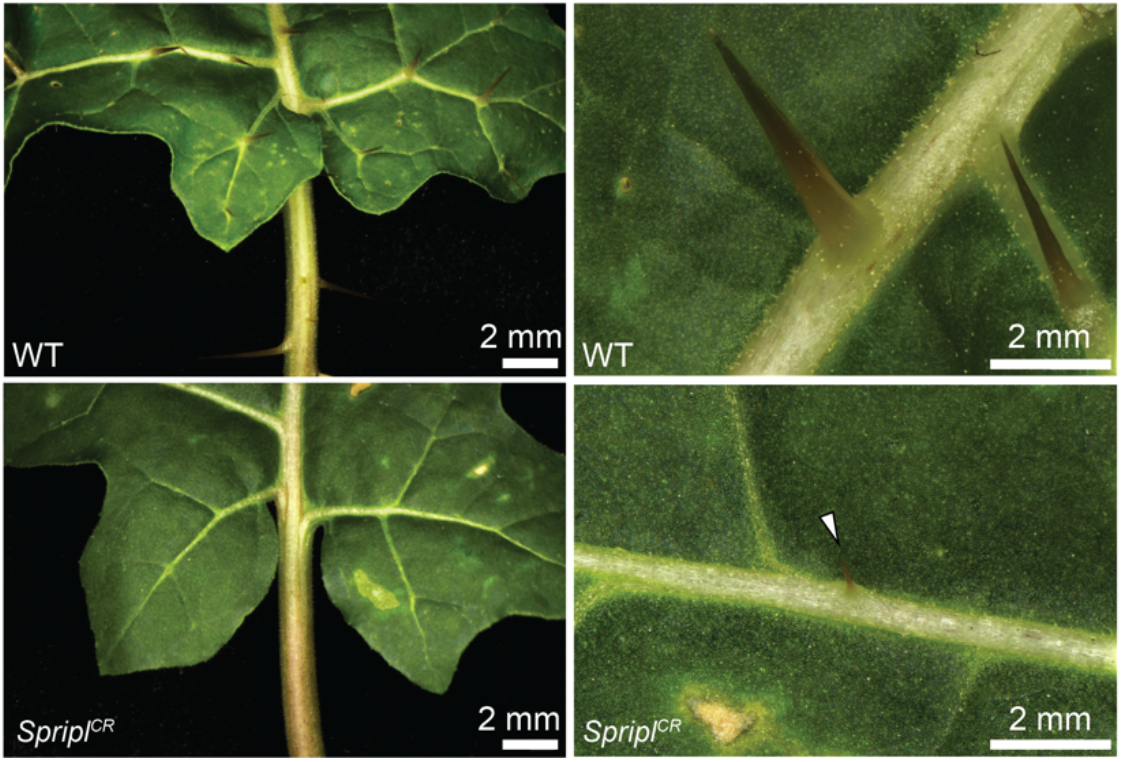
Small prickles are occasionally observed on *pl*^*CR*^ plants. Bright-field images of leaves of Forest nightshade (*S. prinophyllum*) WT and *Spripl*^*CR*^ plants. Arrowhead indicates the presence of a small prickle along a *Spripl*^*CR*^ leaf vein.

## Notes

### Competing Interest Statement

The authors have declared no competing interest.

## References

1. Y. F. Chan, M. E. Marks, F. C. Jones, G. Villarreal, M. D. Shapiro, S. D. Brady, A. M. Southwick, D. M. Absher, J. Grimwood, J. Schmutz, R. M. Myers, D. Petrov, B. Jónsson, D. Schluter, M. A. Bell, D. M. Kingsley, Adaptive Evolution of Pelvic Reduction in Sticklebacks by Recurrent Deletion of a Pitx1 Enhancer. Science 327, 302–305 (2010). doi:10.1126/science.1182213.

2. V. Konečná, S. Bray, J. Vlček, M. Bohutínská, D. Požárová, R. R. Choudhury, A. Bollmann-Giolai, P. Flis, D. E. Salt, C. Parisod, L. Yant, F. Kolář, Parallel adaptation in autopolyploid Arabidopsis arenosa is dominated by repeated recruitment of shared alleles. Nat. Commun. 12, 4979 (2021). doi:10.1038/s41467-021-25256-5.

3. M. Bohutínská, C. L. Peichel, Divergence time shapes gene reuse during repeated adaptation. Trends Ecol. Evol., S0169-5347(23)00325–7 (2023). doi:10.1016/j.tree.2023.11.007.

4. J. Arendt, D. Reznick, Convergence and parallelism reconsidered: what have we learned about the genetics of adaptation? Trends Ecol. Evol. 23, 26–32 (2008). doi:10.1016/j.tree.2007.09.011.

5. T. C. Coverdale, Defence emergence during early ontogeny reveals important differences between spines, thorns and prickles. Ann. Bot. 124, iii–iv (2019). doi:10.1093/aob/mcz189.

6. F. Gallenmüller, A. Feus, K. Fiedler, T. Speck, Rose Prickles and Asparagus Spines – Different Hook Structures as Attachment Devices in Climbing Plants. PLoS ONE 10, e0143850 (2015). doi:10.1371/journal.pone.0143850.

7. Water Worlds, The Green Planet (2022).

8. M.-C. Zhong, X.-D. Jiang, G.-Q. Yang, W.-H. Cui, Z.-Q. Suo, W.-J. Wang, Y.-B. Sun, D. Wang, X.-C. Cheng, X.-M. Li, X. Dong, K.-X. Tang, D.-Z. Li, J.-Y. Hu, Rose without prickle: genomic insights linked to moisture adaptation. Natl. Sci. Rev. 8, nwab092 (2021). doi:10.1093/nsr/nwab092.

9. E. Gagnon, R. Hilgenhof, A. Orejuela, A. McDonnell, G. Sablok, X. Aubriot, L. Giacomin, Y. Gouvêa, T. Bragionis, J. R. Stehmann, L. Bohs, S. Dodsworth, C. Martine, P. Poczai, S. Knapp, T. Särkinen, Phylogenomic discordance suggests polytomies along the backbone of the large genus Solanum. Am. J. Bot. 109, 580–601 (2022). doi:10.1002/ajb2.1827.

10. R. Hilgenhof, E. Gagnon, S. Knapp, X. Aubriot, E. J. Tepe, L. Bohs, L. L. Giacomin, Y. F. Gouvêa, C. T. Martine, A. Orejuela, C. I. Orozco, I. E. Peralta, T. Särkinen, Morphological trait evolution in Solanum (Solanaceae): Evolutionary lability of key taxonomic characters. Taxon 72, 811–847 (2023). doi:10.1002/tax.12990.

11. A. Frary, A. Frary, M.-C. Daunay, K. Huvenaars, R. Mank, S. Doğanlar, QTL hotspots in eggplant (Solanum melongena) detected with a high resolution map and CIM analysis. Euphytica 197, 211–228 (2014). doi:10.1007/s10681-013-1060-6.

12. S. Li, Y. He, D. Li, S. Shi, Y. Wang, X. Tang, H. Ge, Y. Liu, H. Chen, Fine mapping an AUXIN RESPONSE FACTOR, SmARF18, as a candidate gene of the PRICKLE LOCUS that controls prickle absence/presence on various organs in eggplant (Solanum melongena L.). Sci. Hortic. 327, 112874 (2024). doi:10.1016/j.scienta.2024.112874.

13. T. Kurakawa, N. Ueda, M. Maekawa, K. Kobayashi, M. Kojima, Y. Nagato, H. Sakakibara, J. Kyozuka, Direct control of shoot meristem activity by a cytokinin-activating enzyme. Nature 445, 652–655 (2007). doi:10.1038/nature05504.

14. J. G. Monroe, T. Srikant, P. Carbonell-Bejerano, C. Becker, M. Lensink, M. Exposito-Alonso, M. Klein, J. Hildebrandt, M. Neumann, D. Kliebenstein, M.-L. Weng, E. Imbert, J. Ågren, M. T. Rutter, C. B. Fenster, D. Weigel, Mutation bias reflects natural selection in Arabidopsis thaliana. Nature 602, 101–105 (2022). doi:10.1038/s41586-021-04269-6.

15. L. Hua, D. R. Wang, L. Tan, Y. Fu, F. Liu, L. Xiao, Z. Zhu, Q. Fu, X. Sun, P. Gu, H. Cai, S. R. McCouch, C. Sun, LABA1, a Domestication Gene Associated with Long, Barbed Awns in Wild Rice. Plant Cell 27, 1875–1888 (2015). doi:10.1105/tpc.15.00260.

16. S. G. Milner, M. Jost, S. Taketa, E. R. Mazón, A. Himmelbach, M. Oppermann, S. Weise, H. Knüpffer, M. Basterrechea, P. König, D. Schüler, R. Sharma, R. K. Pasam, T. Rutten, G. Guo, D. Xu, J. Zhang, G. Herren, T. Müller, S. G. Krattinger, B. Keller, Y. Jiang, M. Y. González, Y. Zhao, A. Habekuß, S. Färber, F. Ordon, M. Lange, A. Börner, Graner, J. C. Reif, U. Scholz, M. Mascher, N. Stein, Genebank genomics highlights the diversity of a global barley collection. Nat. Genet. 51, 319–326 (2019). doi:10.1038/s41588-018-0266-x.

17. M.-J. Liu, J. Zhao, Q.-L. Cai, G.-C. Liu, J.-R. Wang, Z.-H. Zhao, P. Liu, L. Dai, G. Yan, W.-J. Wang, X.-S. Li, Y. Chen, Y.-D. Sun, Z.-G. Liu, M.-J. Lin, J. Xiao, Y.-Y. Chen, X.-F. Li, B. Wu, Y. Ma, J.-B. Jian, W. Yang, Z. Yuan, X.-C. Sun, Y.-L. Wei, L.-L. Yu, C. Zhang, S.-G. Liao, R.-J. He, X.-M. Guang, Z. Wang, Y.-Y. Zhang, L.-H. Luo, The complex jujube genome provides insights into fruit tree biology. Nat. Commun. 5, 5315 (2014). doi:10.1038/ncomms6315.

18. J. Huang, C. Zhang, X. Zhao, Z. Fei, K. Wan, Z. Zhang, X. Pang, X. Yin, Y. Bai, X. Sun, L. Gao, R. Li, J. Zhang, X. Li, The Jujube Genome Provides Insights into Genome Evolution and the Domestication of Sweetness/Acidity Taste in Fruit Trees. PLoS Genet. 12, e1006433 (2016). doi:10.1371/journal.pgen.1006433.

19. L.-Y. Shen, H. Luo, X.-L. Wang, X.-M. Wang, X.-J. Qiu, H. Liu, S.-S. Zhou, K.-H. Jia, S. Nie, Y.-T. Bao, R.-G. Zhang, Q.-Z. Yun, Y.-H. Chai, J.-Y. Lu, Y. Li, S.-W. Zhao, J.-F. Mao, S.-G. Jia, Y.-M. Mao, Chromosome-Scale Genome Assembly for Chinese Sour Jujube and Insights Into Its Genome Evolution and Domestication Signature. Front. Plant Sci. 12, 773090 (2021). doi:10.3389/fpls.2021.773090.

20. S. Cheng, E. Van Den Bergh, P. Zeng, X. Zhong, J. Xu, X. Liu, J. Hofberger, S. De Bruijn, A. S. Bhide, C. Kuelahoglu, C. Bian, J. Chen, G. Fan, K. Kaufmann, J. C. Hall, A. Becker, A. Bräutigam, A. P. M. Weber, C. Shi, Z. Zheng, W. Li, M. Lv, Y. Tao, J. Wang, H. Zou, Z. Quan, J. M. Hibberd, G. Zhang, X.-G. Zhu, X. Xu, M. E. Schranz, The Tarenaya hassleriana Genome Provides Insight into Reproductive Trait and Genome Evolution of Crucifers. Plant Cell 25, 2813–2830 (2013). doi:10.1105/tpc.113.113480.

21. P. K. Latz, Bushfires & Bushtucker: Aboriginal Plant Use in Central Australia (IAD Press, Alice Springs, Australia, 1995).

22. J. Lee, M. Shah, S. Ballouz, M. Crow, J. Gillis, CoCoCoNet: conserved and comparative co-expression across a diverse set of species. Nucleic Acids Res. 48, W566–W571 (2020). doi:10.1093/nar/gkaa348.

23. Y. Han, Y. Jiao, APETALA1 establishes determinate floral meristem through regulating cytokinins homeostasis in Arabidopsis. Plant Signal. Behav. 10, e989039 (2015). doi:10.4161/15592324.2014.989039.

24. C. Brooks, V. Nekrasov, Z. B. Lippman, J. Van Eck, Efficient Gene Editing in Tomato in the First Generation Using the Clustered Regularly Interspaced Short Palindromic Repeats/CRISPR-Associated9 System. Plant Phys. 166, 1292–1297 (2014). doi:10.1104/pp.114.247577.

25. A. Martin, V. Orgogozo, The Loci of Repeated Evolution: A Catalog of Genetic Hotspots of Phenotypic Variation. Evolution 67, 1235–1250 (2013). doi:10.1111/evo.12081.

26. J. Parker, G. Tsagkogeorga, J. A. Cotton, Y. Liu, P. Provero, E. Stupka, S. J. Rossiter, Genome-wide signatures of convergent evolution in echolocating mammals. Nature 502, 228–231 (2013). doi:10.1038/nature12511.

27. A. D. Foote, Y. Liu, G. W. C. Thomas, T. Vinař, J. Alföldi, J. Deng, S. Dugan, C. E. Van Elk, M. E. Hunter, V. Joshi, Z. Khan, C. Kovar, S. L. Lee, K. Lindblad-Toh, A. Mancia, R. Nielsen, X. Qin, J. Qu, B. J. Raney, N. Vijay, J. B. W. Wolf, M. W. Hahn, D. M. Muzny, K. C. Worley, M. T. P. Gilbert, R. A. Gibbs, Convergent evolution of the genomes of marine mammals. Nat. Genet. 47, 272–275 (2015). doi:10.1038/ng.3198.

28. G. Chomicki, G. Burin, L. Busta, J. Gozdzik, R. Jetter, B. Mortimer, U. Bauer, Convergence in carnivorous pitcher plants reveals a mechanism for composite trait evolution. Science 383, 108–113 (2024). doi:10.1126/science.ade0529.

29. G. Wistow, Lens crystallins: gene recruitment and evolutionary dynamism. Trends Biochem. Sci. 18, 301–306 (1993). doi:10.1016/0968-0004(93)90041-K.

30. N. Shubin, C. Tabin, S. Carroll, Deep homology and the origins of evolutionary novelty. Nature 457, 818–823 (2009). doi:10.1038/nature07891.

31. D. L. Stern, The genetic causes of convergent evolution. Nat. Rev. Genet. 14, 751–764 (2013). doi:10.1038/nrg3483.

32. P. N. Lee, P. Callaerts, H. G. De Couet, M. Q. Martindale, Cephalopod Hox genes and the origin of morphological novelties. Nature 424, 1061–1065 (2003). doi:10.1038/nature01872.

33. T. Kuroha, H. Tokunaga, M. Kojima, N. Ueda, T. Ishida, S. Nagawa, H. Fukuda, K. Sugimoto, H. Sakakibara, Functional Analyses of LONELY GUY Cytokinin-Activating Enzymes Reveal the Importance of the Direct Activation Pathway in Arabidopsis. Plant Cell 21, 3152–3169 (2009). doi:10.1105/tpc.109.068676.

34. L. Chen, G. B. Jameson, Y. Guo, J. Song, P. E. Jameson, The LONELY GUY gene family: from mosses to wheat, the key to the formation of active cytokinins in plants. Plant Biotechnol. J. 20, 625–645 (2022). doi:10.1111/pbi.13783.

35. A. M. Rashotte, The evolution of cytokinin signaling and its role in development before Angiosperms. Semin. Cell Biol. 109, 31–38 (2021). doi:10.1016/j.semcdb.2020.06.010.

36. T. Eviatar-Ribak, A. Shalit-Kaneh, L. Chappell-Maor, Z. Amsellem, Y. Eshed, E. Lifschitz, A Cytokinin-Activating Enzyme Promotes Tuber Formation in Tomato. Curr. Biol. 23, 1057–1064 (2013). doi:10.1016/j.cub.2013.04.061.

37. L. Tirichine, N. Sandal, L. H. Madsen, S. Radutoiu, S. Albrektsen, S. Sato, E. Asamizu, S. Tabata, J. Stougaard, A Gain-of-Function Mutation in a Cytokinin Receptor Triggers Spontaneous Root Nodule Organogenesis. Science 315, 104–107 (2007). doi:10.1126/science.1132397.

38. V. Irish, The ABC model of floral development. Curr. Biol. 27, R887–R890 (2017). doi:10.1016/j.cub.2017.03.045.

39. T. Kudo, T. Kiba, H. Sakakibara, Metabolism and long-distance translocation of cytokinins. J. Integr. Plant Biol. 52, 53–60 (2010). doi:10.1111/j.1744-7909.2010.00898.x.

40. R. H. G. Ranil, H. M. L. Niran, M. Plazas, R. M. Fonseka, H. H. Fonseka, S. Vilanova, I. Andújar, P. Gramazio, A. Fita, J. Prohens, Improving seed germination of the eggplant rootstock Solanum torvum by testing multiple factors using an orthogonal array design. Sci. Hortic. 193, 174–181 (2015). doi:10.1016/j.scienta.2015.07.030.

41. A. M. McClung, J. D. Edwards, M. H. Jia, T. D. Huggins, H. E. Bockelman, M. L. Ali, G. C. Eizenga, Enhancing the searchability, breeding utility, and efficient management of germplasm accessions in the USDA−ARS rice collection. Crop Sci. 60, 3191–3211 (2020). doi:10.1002/csc2.20256.

42. B. Kouassi, J. Prohens, P. Gramazio, A. B. Kouassi, S. Vilanova, A. Galán-Ávila, F. J. Herraiz, A. Kouassi, J. M. Seguí-Simarro, M. Plazas, Development of backcross generations and new interspecific hybrid combinations for introgression breeding in eggplant (Solanum melongena). Sci. Hortic. 213, 199–207 (2016). doi:10.1016/j.scienta.2016.10.039.

43. M. Plazas, S. Vilanova, P. Gramazio, A. Rodríguez-Burruezo, A. Fita, F. J. Herraiz, R. Ranil, R. Fonseka, L. Niran, H. Fonseka, B. Kouassi, A. Kouassi, A. Kouassi, J. Prohens, Interspecific Hybridization between Eggplant and Wild Relatives from Different Genepools. J. Amer. Soc. Hort. Sci. 141, 34–44 (2016). doi:10.21273/JASHS.141.1.34.

44. M. Plazas, P. Gramazio, S. Vilanova, A. B. Kouassi, R. M. Fonseka, M. Rakha, E. Garcia-Fortea, G. Mangino, K. B. A. Kouassi, H. Fonseka, D. Taher, A. Kouassi, G. Villanueva, A. Arrones, D. Alonso, J. Prohens, Introgression breeding from crop wild relatives in eggplant landraces for adaptation to climate change. Crop wild relative, 32–36 (2020).

45. L. Barchi, A. Acquadro, D. Alonso, G. Aprea, L. Bassolino, O. Demurtas, P. Ferrante, P. Gramazio, P. Mini, E. Portis, D. Scaglione, L. Toppino, S. Vilanova, M. J. Díez, G. L. Rotino, S. Lanteri, J. Prohens, G. Giuliano, Single Primer Enrichment Technology (SPET) for High-Throughput Genotyping in Tomato and Eggplant Germplasm. Front. Plant Sci. 10, 1005 (2019). doi:10.3389/fpls.2019.01005.

46. P. Gramazio, H. Yan, T. Hasing, S. Vilanova, J. Prohens, A. Bombarely, Whole-Genome Resequencing of Seven Eggplant (Solanum melongena) and One Wild Relative (S. incanum) Accessions Provides New Insights and Breeding Tools for Eggplant Enhancement. Front. Plant Sci. 10, 1220 (2019). doi:10.3389/fpls.2019.01220.

47. S. Vilanova, D. Alonso, P. Gramazio, M. Plazas, E. García-Fortea, P. Ferrante, M. Schmidt, M. J. Díez, B. Usadel, G. Giuliano, J. Prohens, SILEX: a fast and inexpensive high-quality DNA extraction method suitable for multiple sequencing platforms and recalcitrant plant species. Plant Methods 16, 110 (2020). doi:10.1186/s13007-020-00652-y.

48. L. Barchi, M. Pietrella, L. Venturini, A. Minio, L. Toppino, A. Acquadro, G. Andolfo, G. Aprea, C. Avanzato, L. Bassolino, C. Comino, A. D. Molin, A. Ferrarini, L. C. Maor, E. Portis, S. Reyes-Chin-Wo, R. Rinaldi, T. Sala, D. Scaglione, P. Sonawane, P. Tononi, E. Almekias-Siegl, E. Zago, M. R. Ercolano, A. Aharoni, M. Delledonne, G. Giuliano, S. Lanteri, G. L. Rotino, A chromosome-anchored eggplant genome sequence reveals key events in Solanaceae evolution. Sci. Rep. 9, 11769 (2019). doi:10.1038/s41598-019-47985-w.

49. C. T. Wittwer, G. H. Reed, C. N. Gundry, J. G. Vandersteen, R. J. Pryor, High-Resolution Genotyping by Amplicon Melting Analysis Using LCGreen. Clin. Chem. 49, 853–860 (2003). doi:10.1373/49.6.853.

50. M. Kanakachari, A. U. Solanke, N. Prabhakaran, I. Ahmad, G. Dhandapani, N. Jayabalan, P. A. Kumar, Evaluation of Suitable Reference Genes for Normalization of qPCR Gene Expression Studies in Brinjal (Solanum melongena L.) During Fruit Developmental Stages. Appl Biochem Biotechnol 178, 433–450 (2016). doi:10.1007/s12010-015-1884-8.

51. J. J. Doyle, J. L. Doyle, A rapid DNA isolation procedure for small quantities of fresh leaf tissue. Phytochem. Bull. 19, 11–15 (1987).

52. H. Takagi, A. Abe, K. Yoshida, S. Kosugi, S. Natsume, C. Mitsuoka, A. Uemura, H. Utsushi, M. Tamiru, S. Takuno, H. Innan, L. M. Cano, S. Kamoun, R. Terauchi, QTL-seq: rapid mapping of quantitative trait loci in rice by whole genome resequencing of DNA from two bulked populations. Plant J. 74, 174–183 (2013). doi:10.1111/tpj.12105.

53. M. Alonge, X. Wang, M. Benoit, S. Soyk, L. Pereira, L. Zhang, H. Suresh, S. Ramakrishnan, F. Maumus, D. Ciren, Y. Levy, T. H. Harel, G. Shalev-Schlosser, Z. Amsellem, H. Razifard, A. L. Caicedo, D. M. Tieman, H. Klee, M. Kirsche, S. Aganezov, T. R. Ranallo-Benavidez, Z. H. Lemmon, J. Kim, G. Robitaille, M. Kramer, S. Goodwin, W. R. McCombie, S. Hutton, J. Van Eck, J. Gillis, Y. Eshed, F. J. Sedlazeck, E. Van Der Knaap, M. C. Schatz, Z. B. Lippman, Major Impacts of Widespread Structural Variation on Gene Expression and Crop Improvement in Tomato. Cell 182, 145-161.e23 (2020). doi:10.1016/j.cell.2020.05.021.

54. T. R. Ranallo-Benavidez, K. S. Jaron, M. C. Schatz, GenomeScope 2.0 and Smudgeplot for reference-free profiling of polyploid genomes. Nat. Commun. 11, 1432 (2020). doi:10.1038/s41467-020-14998-3.

55. H. Cheng, G. T. Concepcion, X. Feng, H. Zhang, H. Li, Haplotype-resolved de novo assembly using phased assembly graphs with hifiasm. Nat. Methods 18, 170–175 (2021). doi:10.1038/s41592-020-01056-5.

56. M. Alonge, L. Lebeigle, M. Kirsche, K. Jenike, S. Ou, S. Aganezov, X. Wang, Z. B. Lippman, M. C. Schatz, S. Soyk, Automated assembly scaffolding using RagTag elevates a new tomato system for high-throughput genome editing. Genome Biol. 23, 258 (2022). doi:10.1186/s13059-022-02823-7.

57. A. Rhie, B. P. Walenz, S. Koren, A. M. Phillippy, Merqury: reference-free quality, completeness, and phasing assessment for genome assemblies. Genome Biol. 21, 245 (2020). doi:10.1186/s13059-020-02134-9.

58. A. Dobin, C. A. Davis, F. Schlesinger, J. Drenkow, C. Zaleski, S. Jha, P. Batut, M. Chaisson, T. R. Gingeras, STAR: ultrafast universal RNA-seq aligner. Bioinformatics 29, 15–21 (2013). doi:10.1093/bioinformatics/bts635.

59. S. Kovaka, A. V. Zimin, G. M. Pertea, R. Razaghi, S. L. Salzberg, M. Pertea, Transcriptome assembly from long-read RNA-seq alignments with StringTie2. Genome Biol 20, 278 (2019). doi:10.1186/s13059-019-1910-1.

60. D. Mapleson, L. Venturini, G. Kaithakottil, D. Swarbreck, Efficient and accurate detection of splice junctions from RNA-seq with Portcullis. GigaScience 7 (2018). doi:10.1093/gigascience/giy131.

61. P. S. Hosmani, M. Flores-Gonzalez, H. Van De Geest, F. Maumus, L. V. Bakker, E. Schijlen, J. Van Haarst, J. Cordewener, G. Sanchez-Perez, S. Peters, Z. Fei, J. J. Giovannoni, L. A. Mueller, S. Saha, An improved de novo assembly and annotation of the tomato reference genome using single-molecule sequencing, Hi-C proximity ligation and optical maps. bioRxiv (2019). doi:10.1101/767764.

62. L. Barchi, M. T. Rabanus-Wallace, J. Prohens, L. Toppino, S. Padmarasu, E. Portis, G. L. Rotino, N. Stein, S. Lanteri, G. Giuliano, Improved genome assembly and pan-genome provide key insights into eggplant domestication and breeding. Plant J. 107, 579–596 (2021). doi:10.1111/tpj.15313.

63. A. Shumate, S. L. Salzberg, Liftoff: accurate mapping of gene annotations. Bioinformatics 37, 1639–1643 (2021). doi:10.1093/bioinformatics/btaa1016.

64. T. D. Wu, C. K. Watanabe, GMAP: a genomic mapping and alignment program for mRNA and EST sequences. Bioinformatics 21, 1859–1875 (2005). doi:10.1093/bioinformatics/bti310.

65. H. Li, Minimap2: pairwise alignment for nucleotide sequences. Bioinformatics 34, 3094–3100 (2018). doi:10.1093/bioinformatics/bty191.

66. The Potato Genome Sequencing Consortium, Genome sequence and analysis of the tuber crop potato. Nature 475, 189–195 (2011). doi:10.1038/nature10158.

67. L. Venturini, S. Caim, G. G. Kaithakottil, D. L. Mapleson, D. Swarbreck, Leveraging multiple transcriptome assembly methods for improved gene structure annotation. GigaScience 7 (2018). doi:10.1093/gigascience/giy093.

68. A. Kuno, Y. Ikeda, S. Ayabe, K. Kato, K. Sakamoto, S. R. Suzuki, K. Morimoto, A. Wakimoto, N. Mikami, M. Ishida, N. Iki, Y. Hamada, M. Takemura, Y. Daitoku, Y. Tanimoto, T. T. H. Dinh, K. Murata, M. Hamada, M. Muratani, A. Yoshiki, F. Sugiyama, S. Takahashi, S. Mizuno, DAJIN enables multiplex genotyping to simultaneously validate intended and unintended target genome editing outcomes. PLoS Biol 20, e3001507 (2022). doi:10.1371/journal.pbio.3001507.

69. D. M. Emms, S. Kelly, OrthoFinder: phylogenetic orthology inference for comparative genomics. Genome Biol. 20, 238 (2019). doi:10.1186/s13059-019-1832-y.

70. H. Li, Protein-to-genome alignment with miniprot. Bioinformatics 39, btad014 (2023). doi:10.1093/bioinformatics/btad014.

71. A. J. Hart, S. Ginzburg, M. Xu, C. R. Fisher, N. Rahmatpour, J. B. Mitton, R. Paul, J. L. Wegrzyn, ENTAP: Bringing faster and smarter functional annotation to non-model eukaryotic transcriptomes. Mol. Ecol. Resour. 20, 591–604 (2020). doi:10.1111/1755-0998.13106.

72. M. Van Bel, F. Silvestri, E. M. Weitz, L. Kreft, A. Botzki, F. Coppens, K. Vandepoele, PLAZA 5.0: extending the scope and power of comparative and functional genomics in plants. Nucleic Acids Res. 50, D1468–D1474 (2022). doi:10.1093/nar/gkab1024.

73. R. Apweiler, A. Bairoch, C. H. Wu, W. C. Barker, B. Boeckmann, S. Ferro, E. Gasteiger, H. Huang, R. Lopez, M. Magrane, M. J. Martin, D. A. Natale, C. O’Donovan, N. Redaschi, L.-S. L. Yeh, UniProt: the Universal Protein knowledgebase. Nucleic Acids Res. 32, D115–119 (2004). doi:10.1093/nar/gkh131.

74. P. Jones, D. Binns, H.-Y. Chang, M. Fraser, W. Li, C. McAnulla, H. McWilliam, J. Maslen, A. Mitchell, G. Nuka, S. Pesseat, A. F. Quinn, A. Sangrador-Vegas, M. Scheremetjew, S.-Y. Yong, R. Lopez, S. Hunter, InterProScan 5: genome-scale protein function classification. Bioinformatics 30, 1236–1240 (2014). doi:10.1093/bioinformatics/btu031.

75. M. Van Bel, S. Proost, C. Van Neste, D. Deforce, Y. Van de Peer, K. Vandepoele, TRAPID: an efficient online tool for the functional and comparative analysis of de novo RNA-Seq transcriptomes. Genome Biol. 14, R134 (2013). doi:10.1186/gb-2013-14-12-r134.

76. R.-G. Zhang, G.-Y. Li, X.-L. Wang, J. Dainat, Z.-X. Wang, S. Ou, Y. Ma, TEsorter: An accurate and fast method to classify LTR-retrotransposons in plant genomes. Hortic. Res. 9, uhac017 (2022). doi:10.1093/hr/uhac017.

77. M. Manni, M. R. Berkeley, M. Seppey, E. M. Zdobnov, BUSCO: Assessing Genomic Data Quality and Beyond. Curr. Protoc. 1, e323 (2021). doi:10.1002/cpz1.323.

78. T. Särkinen, M. Staats, J. E. Richardson, R. S. Cowan, F. T. Bakker, How to Open the Treasure Chest? Optimising DNA Extraction from Herbarium Specimens. PLoS ONE 7, e43808 (2012). doi:10.1371/journal.pone.0043808.

79. L. D. Shepherd, T. G. B. McLay, Two micro-scale protocols for the isolation of DNA from polysaccharide-rich plant tissue. J. Plant Res. 124, 311–314 (2011). doi:10.1007/s10265-010-0379-5.

80. S. Li, G. Yang, S. Yang, J. Just, H. Yan, N. Zhou, H. Jian, Q. Wang, M. Chen, X. Qiu, H. Zhang, X. Dong, X. Jiang, Y. Sun, M. Zhong, M. Bendahmane, G. Ning, H. Ge, J.-Y. Hu, K. Tang, The development of a high-density genetic map significantly improves the quality of reference genome assemblies for rose. Sci. Rep. 9, 5985 (2019). doi:10.1038/s41598-019-42428-y.

81. M. Stanke, B. Morgenstern, AUGUSTUS: a web server for gene prediction in eukaryotes that allows user-defined constraints. Nucleic Acids Res. 33, W465–467 (2005). doi:10.1093/nar/gki458.

82. G. S. C. Slater, E. Birney, Automated generation of heuristics for biological sequence comparison. BMC Bioinform. 6, 31 (2005). doi:10.1186/1471-2105-6-31.

83. A. M. Bolger, M. Lohse, B. Usadel, Trimmomatic: a flexible trimmer for Illumina sequence data. Bioinformatics 30, 2114–2120 (2014). doi:10.1093/bioinformatics/btu170.

84. K. Katoh, D. M. Standley, MAFFT multiple sequence alignment software version 7: improvements in performance and usability. Mol. Biol. Evol. 30, 772–780 (2013). doi:10.1093/molbev/mst010.

85. A. Stamatakis, T. Ludwig, H. Meier, RAxML-III: a fast program for maximum likelihood-based inference of large phylogenetic trees. Bioinformatics 21, 456–463 (2005). doi:10.1093/bioinformatics/bti191.

86. M. A. Miller, W. Pfeiffer, T. Schwartz, “Creating the CIPRES Science Gateway for inference of large phylogenetic trees” in 2010 Gateway Computing Environments Workshop (GCE) (IEEE, 2010), pp. 1–8.

87. G. Yu, D. K. Smith, H. Zhu, Y. Guan, T. T. Lam, GGTREE : an R package for visualization and annotation of phylogenetic trees with their covariates and other associated data. Methods Ecol. Evol. 8, 28–36 (2017). doi:10.1111/2041-210X.12628.

88. Z. H. Lemmon, S. J. Park, K. Jiang, J. Van Eck, M. C. Schatz, Z. B. Lippman, The evolution of inflorescence diversity in the nightshades and heterochrony during meristem maturation. Genome Res. 26, 1676–1686 (2016). doi:10.1101/gr.207837.116.

89. T. Tamura, M. Nei, Estimation of the number of nucleotide substitutions in the control region of mitochondrial DNA in humans and chimpanzees. Mol. Biol. Evol. 10, 512–526 (1993). doi:10.1093/oxfordjournals.molbev.a040023.

90. J. Van Eck, P. Keen, M. Tjahjadi, “Agrobacterium tumefaciens-Mediated Transformation of Tomato” in Transgenic Plants, S. Kumar, P. Barone, M. Smith, Eds. (Springer New York, New York, NY, 2019)vol. 1864 of Methods in Molecular Biology, pp. 225–234.

91. D. Kuznetsov, F. Tegenfeldt, M. Manni, M. Seppey, M. Berkeley, E. V. Kriventseva, E. M. Zdobnov, OrthoDB v11: annotation of orthologs in the widest sampling of organismal diversity. Nucleic Acids Res. 51, D445–D451 (2023). doi:10.1093/nar/gkac998.

92. M. Crow, H. Suresh, J. Lee, J. Gillis, Coexpression reveals conserved gene programs that co-vary with cell type across kingdoms. Nucleic Acids Res. 50, 4302–4314 (2022). doi:10.1093/nar/gkac276.

